# Membrane protein condensates polymerize actin and form filopodia

**DOI:** 10.1101/2025.10.09.681077

**Authors:** Shannon Bareesel, Thomas J. Böddeker, Willem Bintig, Domonkos Nagy-Herczeg, Max Ruwolt, Vasiliki Syropoulou, Cristina Kroon, Patricia Kreis, Susanne Wegmann, Michal Szczepek, Christian M.T. Spahn, Patrick Scheerer, George Leondaritis, Fan Liu, Roland L. Knorr, Britta J. Eickholt

**Affiliations:** Institute of Molecular Biology and Biochemistry, Charité – Universitätsmedizin Berlin; Berlin, 10117, Germany; Institute of Medical Physics and Biophysics, Group Structural Biology of Cellular Signaling, Charité – Universitätsmedizin Berlin; Berlin, 10117, Germany; Institute of Biology, Humboldt University of Berlin; Berlin, 10115, Germany; Centre for Cancer Cell Reprogramming, Faculty of Medicine, University of Oslo; Oslo, 0379, Norway; Department of Structural Biology, Leibniz Forschungsinstitut für Molekulare Pharmakologie (FMP); Berlin, 13125, Germany; German Center for Neurodegenerative Diseases (DZNE); Berlin, 10117, Germany; Institute of Medical Physics and Biophysics, Charité – Universitätsmedizin Berlin; Berlin, 10117, Germany; Department of Pharmacology, Faculty of Medicine, School of Health Sciences, University of Ioannina; Ioannina, 45110, Greece; Institute of Biosciences, University Research Center Ioannina, University of Ioannina; Ioannina, 45110, Greece; Center for Biochemistry, Faculty of Medicine, University of Cologne; Cologne, 50931, Germany; Graduate School and Faculty of Medicine, The University of Tokyo; Tokyo, 113-0033, Japan; Cologne Excellence Cluster on Aging and Aging-Associated Diseases (CECAD), Faculty of Medicine, University of Cologne; Cologne, 50931, Germany

**Author notes:** These authors contributed equally to this work (Shannon Bareesel, Thomas J. Böddeker).

**Keywords:** PLPPR3, actin polymerization, condensates, liquid-liquid phase separation, neuron growth and branching, filopodia

## Abstract

Neuronal morphogenesis is guided by filopodia, dynamically generated plasma membrane protrusions filled with parallel actin filaments. However, how filopodial actin filaments are locally produced, organized, and maintained remains unclear. The transmembrane protein PLPPR3 induces filopodia in neurons and other cells. We find that the intracellular domain (ICD) of PLPPR3 forms liquid condensates, which exhibit strong co-partitioning of actin monomers. These condensates promote actin polymerization within the condensates at the expense of actin monomers in the environment, consistent with thermodynamic coupling of actin partitioning and polymerization, which we recapitulate in a modified polymerization kinetics model. This mechanism requires favorable actin partitioning into the condensate relative to the environment. Using crosslinking mass spectrometry, we identify a WH2-like actin-affinity domain within the PLPPR3 ICD. Deleting this domain lowers actin partitioning *in vitro* and decreases filopodia formation *in vivo*. Our findings establish a previously unrecognized mechanism for actin network remodeling, in which condensates act as actin sinks, locally boosting monomer concentrations and facilitating polymerization of actin filaments.

**One sentence summary:** Our study uncovers PLPPR3 condensates to locally enrich actin monomers and promote actin polymerization, driving actin network remodeling and neuronal filopodia formation.

## Main Text

Filopodia are slender, finger-like plasma membrane protrusions important for cell migration and sensing. In neuronal cells, filopodia can also mature into specialized compartments including neurites, axon branches and synapses (*1*). Consequently, these structures are particularly important for the formation of neuronal circuits during brain development. Primarily composed of parallel bundles of actin filaments, filopodia formation, stabilization, and dynamics are regulated by various actin-binding proteins (ABPs) and signaling cascades associated with both actin filaments and the plasma membrane (*2*). Among the few membrane proteins implicated in filopodia formation, phospholipid phosphatase-related protein 3 (PLPPR3) has emerged as a regulator of this process in neurons. PLPPR3 is composed of six transmembrane domains and a long intracellular domain (ICD), which is particularly important for filopodia formation (*3*).

### PLPPR3 ICD forms condensates in cells

Previous live-cell imaging of tagged PLPPR3 in hippocampal neurons revealed regularly spaced PLPPR3 puncta along the axonal membrane, marking regions along the axon shaft where actin patches were formed and filopodia preferentially extended (Fig. 1A) (*4*). Interestingly, puncta of endogenous PLPPR3 in hippocampal axons, labelled by antibodies binding the PLPPR3 ICD, can be disrupted by 1,6-hexanediol, a compound known to interfere with many biomolecular condensates (*5, 6*), while washout of the compound recovered punctate PLPPR3 distributions along the axon shaft (Fig. 1B). This suggests that neuronal PLPPR3 puncta represent biomolecular condensates formed by liquid-liquid phase separation (LLPS).

**Fig. 1.**
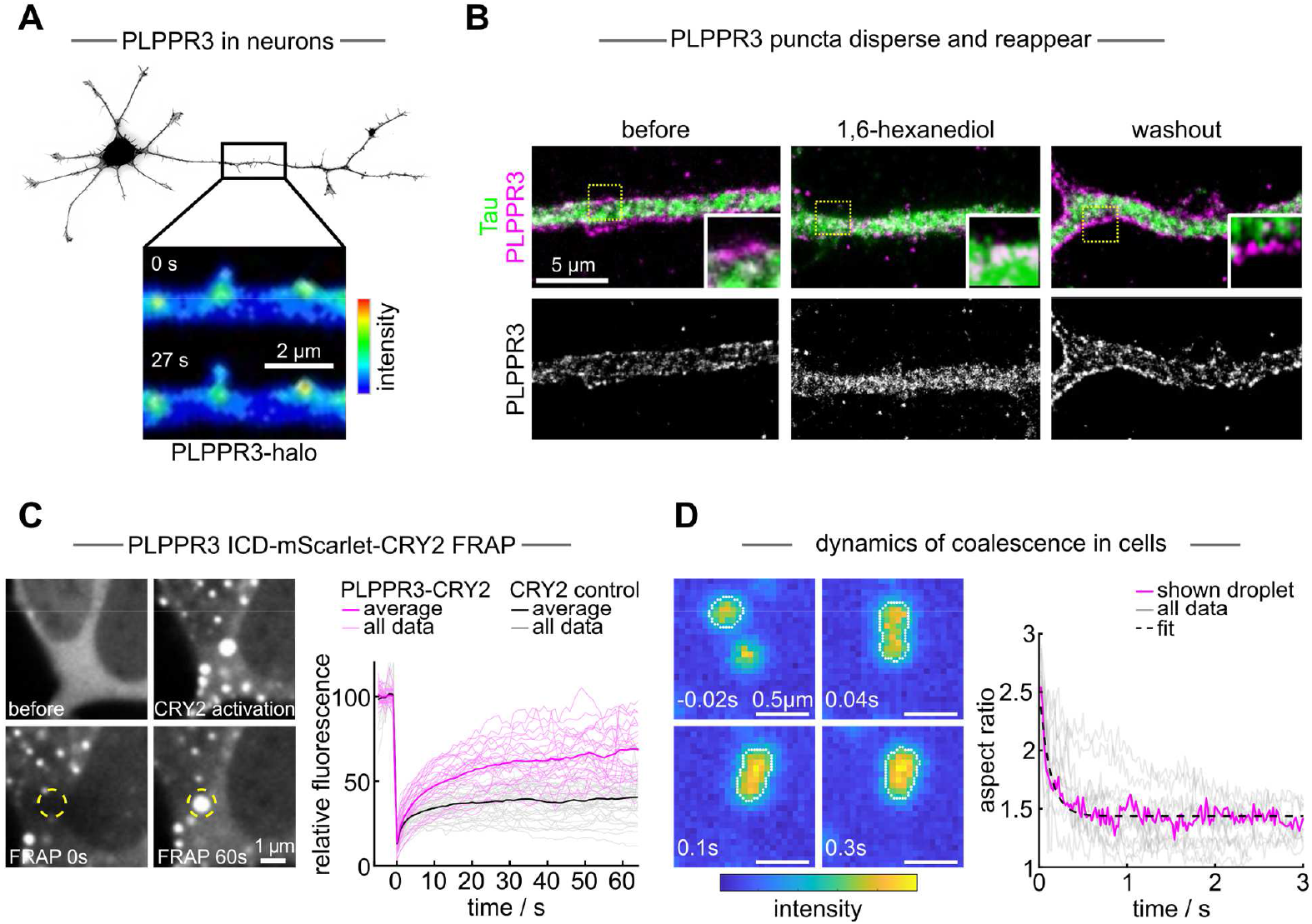
PLPPR3 forms condensates in cells. **(A)** Overexpression of PLPPR3-Halo forms membrane bound puncta in neurons that co-localize with membrane protrusions, see Brosig et al (4). **(B)** PLPPR3 forms puncta along the plasma membrane in axons of hippocampal neurons. Upon 1,6-hexanediol treatment, puncta disperse, whereas puncta reform after washout of hexanediol. **(C)** PLPPR3 ICD-CRY2 puncta recover after photobleaching. Time series of a representative cell before and after CRY2 activation, as well as upon photobleaching and after 60 s (left panel). Average recovery dynamics of N=26 experiments from 7 cells with PLPPR3 ICD-CRY2 and N=39 with CRY2 control from 11 cells. Bold lines show the average with individual experiments shown in the background (right panel). **(D)** Timelapse of two coalescing PLPPR3 ICD-CRY2 condensates. Images show raw data, white dots indicate the analyzed condensate outline (left panel). Aspect ratio over time of the shown coalescence event (magenta) and corresponding exponential fit (dashed line). Remaining data are shown in the background for a total of N=13 coalescence events from 8 cells (right panel).

In neurons, native PLPPR3 puncta are membrane-bound and very small, with an approximately round shape of a diameter of ∼200 nm (Fig. 1B), which is close to the diffraction limit, posing challenges for direct examination of LLPS behavior. For an experimentally accessible approach, we generated optically-inducible cytosolic PLPPR3 condensates by fusing PLPPR3 ICD to CRY2, a blue-light photoreceptor that oligomerizes upon blue-light stimulation (*7*). In HEK293T cells expressing PLPPR3 ICD-mScarlet-CRY2, exposure to blue light induced rapid formation of rounded puncta (Fig. 1C). Cells expressing mScarlet-CRY2 lacking the PLPPR3 ICD also formed puncta upon blue-light exposure, however, these puncta exhibited markedly reduced circularity suggesting more solid-like properties (Supplementary Fig. S1A).

We interrogated the material state of PLPPR3 ICD puncta with fluorescence recovery after photobleaching (FRAP) and bleached the entire puncta to probe for molecular exchange with the cytosol as expected for liquid condensates (Fig. 1C). ICD-mScarlet-CRY2 puncta recovered their fluorescence, reaching about 65% of the initial intensity within 1 min after bleaching. In contrast, control mScarlet-CRY2 puncta exhibited a small degree of recovery within the first few seconds following bleaching, likely attributed to diffusion in the cytosol in the immediate vicinity. These puncta did not recover their fluorescence intensity over longer time, suggesting that the presence of PLPPR3 ICD leads to a higher degree of exchange with the environment.

PLPPR3 ICD-mScarlet-CRY2 puncta also coalesced upon contact, resulting in an increase in size and a decrease in number of puncta over time (Supplementary Fig. S1B). Observing such coalescence events at high frame rates revealed coalescence behavior characteristic of liquid droplets (Fig. 1D). The rate at which droplets coalesce is defined by the inverse capillary velocity, i.e. the ratio of surface tension to viscosity. For PLPPR3 ICD-mScarlet-CRY2 puncta, we measured an average inverse capillary velocity of 1.29 ± 0.77 s/µm (mean±SD, SEM = 0.22 s/µm). After coalescence, puncta typically did not relax into a perfect sphere but retained an elliptical shape with an average aspect ratio of ⟨η/γ⟩ = 1.39 ± 0.21, suggesting that puncta are affected by the surrounding cytosol, as previously observed for other cytosolic protein condensates of similar size (*8, 9*). Overall, these findings confirm that PLPPR3 ICD puncta exhibit liquid-like characteristics and can therefore be classified as condensates.

### PLPPR3 ICD forms liquid-like condensates in vitro

To further characterize PLPPR3 condensates, we purified PLPPR3 ICD from Expi293F cells (Supplementary Fig. S2; Materials and Methods). Structural analysis using circular dichroism spectroscopy suggests that purified PLPPR3 ICD is predominantly unstructured (Supplementary Fig. S3), confirming the structure prediction analysis carried out in PONDR (Fig. 2A). Mapping out a phenomenological phase diagram of PLPPR3 ICD (3% of PLPPR3 ICD labelled with Alexa-488) by systematically varying the concentrations of PLPPR3 ICD and polyethylene glycol (PEG) 8000 – a molecular crowder widely used in condensate studies (*10*) – we found that PLPPR3 ICD formed condensates at a threshold concentration of ∼10 µM at 5% (w/v) PEG (Fig. 2B). Reducing the PEG concentration to 1% raised the threshold concentration to ∼20 µ M. In the absence of PEG, PLPPR3 ICD did not undergo LLPS, suggesting that some degree of molecular crowding is required for condensate formation. We conclude that PLPPR3 ICD has an intrinsic capacity for LLPS.

**Fig. 2.**
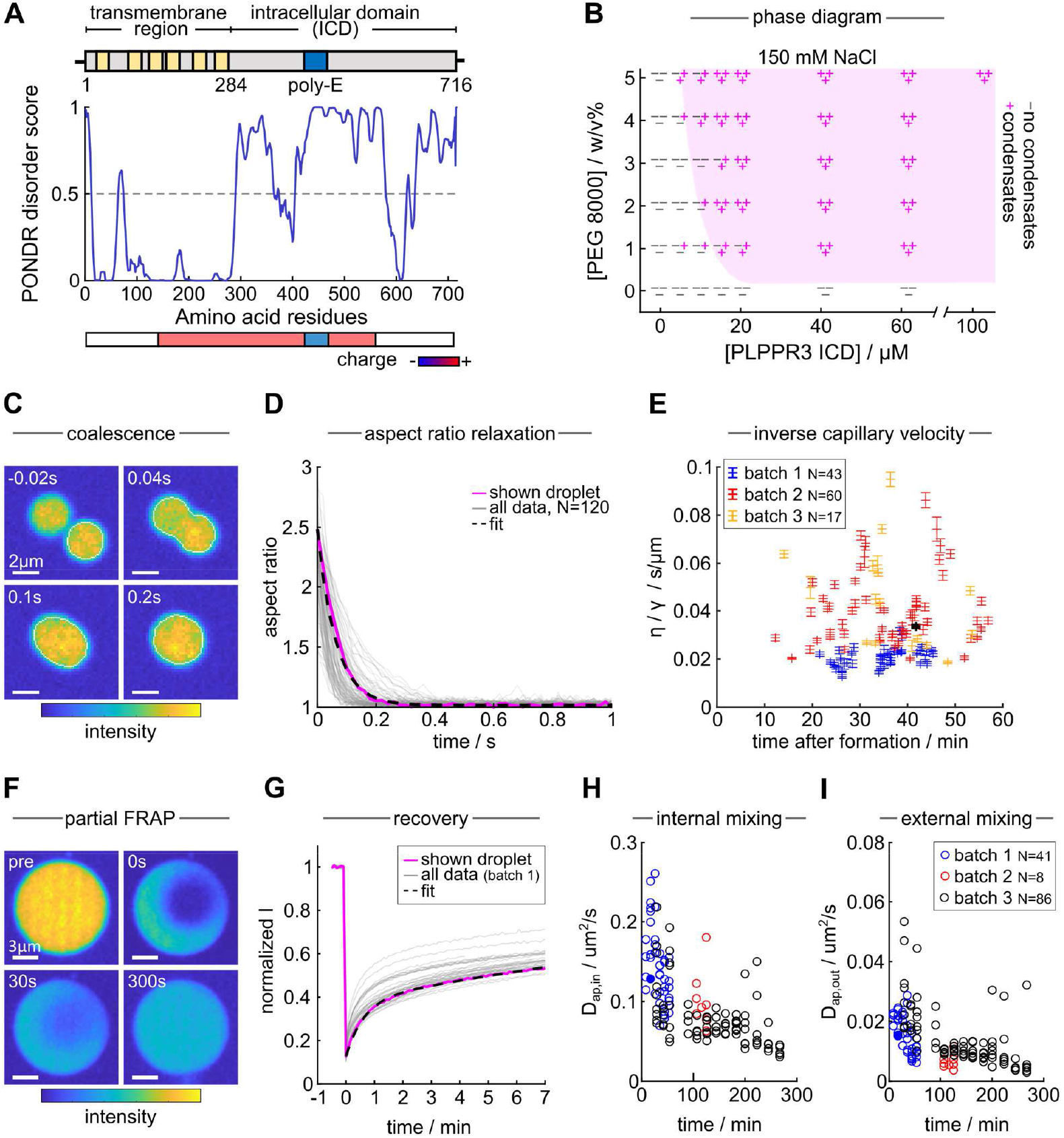
PLPPR3 ICD forms liquid condensates *in vitro*. **(A)** Structural information of PLPPR3. PONDR disorder prediction of PLPPR3 reveals a highly unstructured and partially charged intracellular domain (ICD). **(B)** Phase diagram of PLPPR3 at 150 mM NaCl in 20 mM HEPES buffer under varying PEG concentrations. **(C)** Representative coalescence event of ICD condensates. **(D)** Relaxation of the aspect ratio of all condensates. The condensate shown in **C** is shown in magenta and the exponential fit is shown by the dashed line. **(E)** The inverse capillary velocity *T*/*Y* across three batches of purified PLPPR3 ICD as a function of time after condensate formation by addition of PEG. Measurement of condensate from panel **C** in black. Note that 1 mM ATP was added in experiments with batch 3. **(F)** Representative example of a partial FRAP experiment revealing fast internal mixing and slower exchange with the environment. **(G)** Fluorescence intensity recovery throughout experiments from batch 1 (N=41). The condensate shown in **F** is highlighted in magenta The dashed line shows a double-exponential fit. **(H)** Apparent internal diffusion constant D_ap,in_ as a function of time after induction of phase separation for three batches of protein. **(I)** Apparent external diffusion constant D_ap,out_ for the same data. Measurement from panel **F** shown with filled circle.

To access the material properties of PLPPR3 ICD condensates, we analyzed their coalescence behavior *in vitro* through timelapse imaging. Using non-wetting conditions (contact angle of ∼180° on polyvinyl alcohol coated glass coverslips, see Supplementary Fig. S4), we observed rapid relaxation of the aspect ratio of coalescing condensates to a value of one within about 0.2 s (Fig. 2C, D). Fitting an exponential to the relaxation curves of N=120 coalescence events, we measured an average inverse capillary velocity of ⟨η/γ⟩= 0.034 ± 0.017 s/µm (mean±SD). The relaxation kinetics were thus about 40 times faster than within cells, suggesting a lower surface tension and/or higher viscosity *in vivo* compared to *in vitro*.

We performed these measurements for PLPPR3 protein purified from three batches. While ⟨η/γ⟩differed between samples, we found higher variance within experiments, suggesting a greater impact of experimental conditions, e.g. local coating quality. These results suggest that material properties are overall consistent and independent of time after phase separation triggered by addition of PEG within 1 hour (Fig. 2E).

Next, we performed FRAP measurements on PLPPR3 ICD condensates *in vitro*. Partial FRAP of PLPPR3 ICD condensates identified two distinct recovery regimes: an initial rapid internal mixing within the condensate, followed by a slower recovery phase likely involving exchange with the environment (Fig. 2F). To approximate the observed different time scales, we fitted two exponentials to our data. While this simplified model only estimates the molecular dynamics, it yielded good agreement with the data (Fig. 2G) and enabled us to extract apparent diffusion constants for the fast internal mixing, *D*_*ap,in*_, and slower exchange with the environment, *D*_*ap,out*_. Both *D*_*ap,in*_ and *D*_*ap,out*_ decreased with time after induction of phase separation by addition of PEG, indicative of some degree of ageing (Fig. 2H-I). *D*_*ap,in*_ dropped from ∼0.15 µm^2^/s shortly after phase separation to a value of ∼0.05 µm^2^/s after 4 hours, whereas *D*_*ap,out*_ was about 5-fold smaller at all times. For comparison, the diffusion coefficient of long-chain silicone oil is on the same order of magnitude as *D*_*ap,in*_ of the ICD (*11*). In summary, these results demonstrate that PLPPR3 ICD robustly undergoes LLPS to form dynamic, liquid condensates *in vitro*.

### PLPPR3 ICD condensates localize and enhance actin polymerization

We hypothesize that PLPPR3 ICD regulates filopodia formation by localizing and enhancing actin polymerization through condensate formation. To test this, we mixed 1.2 µ M G-actin (31% fluorescently labeled actin) with PLPPR3 ICD in actin polymerization buffer, in the absence or presence of PEG. In the absence of PEG, this resulted in a homogenous mixture of both G-actin and PLPPR3 ICD. In contrast, when 5% PEG was added to induce LLPS of PLPPR3 ICD, actin partitioned into PLPPR3 ICD condensates (Fig. 3A).

**Fig. 3.**
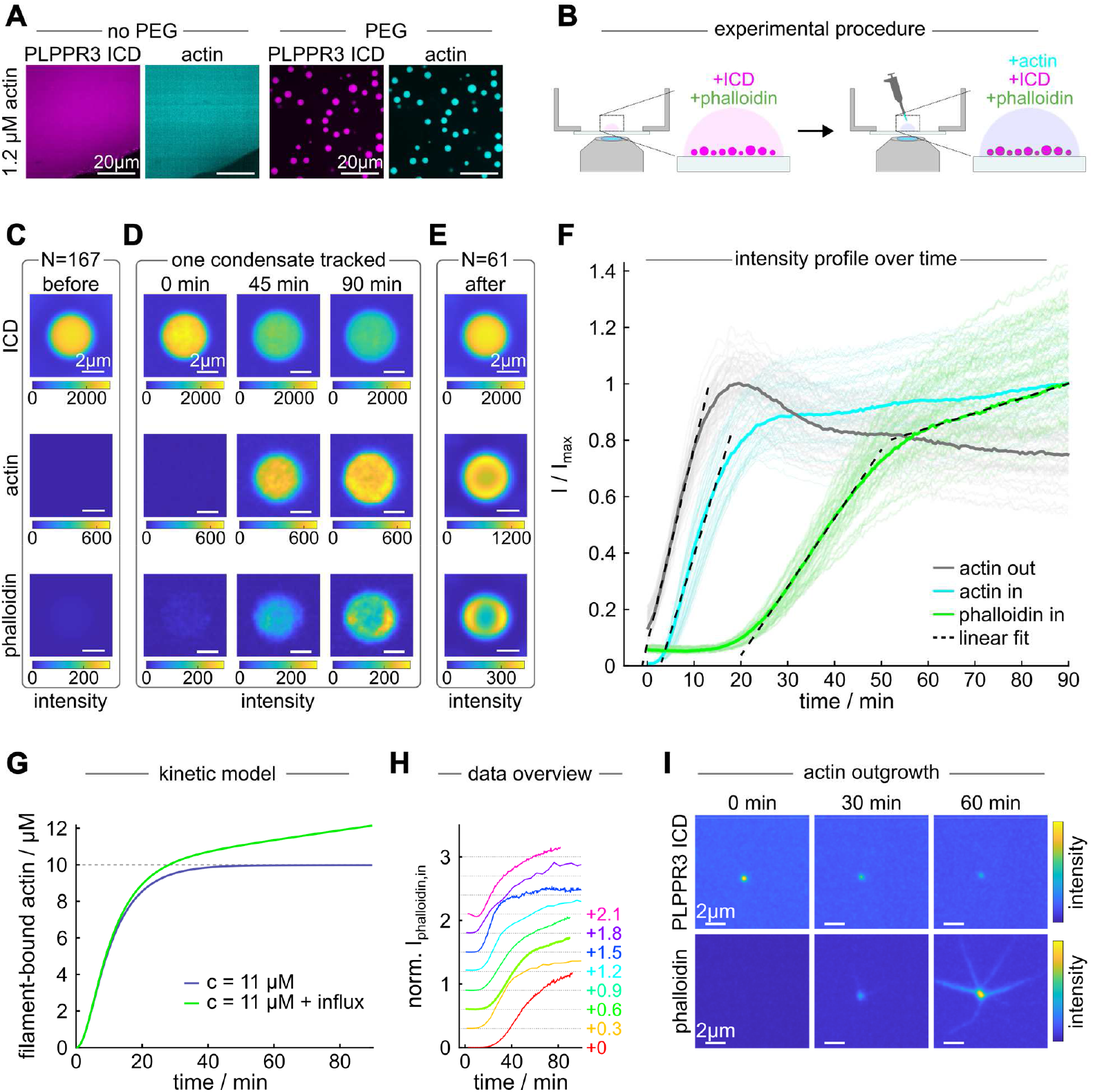
PLPPR3 ICD condensates localize and enhance actin polymerization. **(A)** PLPPR3 intracellular domain (ICD) (20 µM) and actin (1.2 µM) form homogeneous mixtures in the absence of PEG. With PEG, actin strongly partitions into ICD condensates. **(B)** Experimental procedure of the actin polymerization assay. **(C)** Average images of the ICD, actin and phalloidin (F-actin) channel averaged over N = 167 condensates with a radius of 1.98 µm before addition of actin. **(D)** Snapshots of the intensity evolution of one condensate tracked throughout the experiment (Supplementary Movie 1). **(E)** Average images of N = 61 condensates with radius 1.98 µm throughout the sample chamber after acquisition of the time series. Note that the intensity scale for actin and phalloidin is readjusted compared to panel **C** and **D**. (**F**) Intensity profile of actin outside the condensates (gray), inside the condensates (cyan) and phalloidin inside the condensate (green). The highlighted lines are the average over all condensates within the field of view. The transparent lines show the respective intensity profiles of N = 49 tracked condensates. The dashed lines indicate linear fits to the different regimes and define the times of actin arriving in the supernatant (*T*_*act,out*_ = −1.2 ± 0.1 min), actin concentrating in the condensate (*T*_*act,in*_ = 2.9 ± 0.1 min) and the onset of F-actin polymerization (*T*_*phal,in*_ = 20.8 ± 0.1 min), with the start of timelapse imaging at *T* = 0 min. **(G)** The secondary linear regime in F-actin polymerization can be reproduced in a theoretical model of polymerization kinetics by adding a small influx of G-actin. **(H)** Juxtaposition of intensity profiles of the phalloidin channel normalized by intensity after 70 min from 8 independent experiments. An increasing offset is added to the data for better visualization (right side of the panel). Extended data in Supplementary Fig. S7. **(I)** Small ICD condensates, with a size close to the resolution limit, fail to retain growing F-actin.

Next, we investigated actin polymerization within PLPPR3 ICD condensates in real time (Fig. 3B). In these experiments, ICD condensates were formed in actin polymerization buffer containing phalloidin, a highly selective peptide used for visualizing F-actin (*12, 13*), and allowed to settle for at least 60 min. Figure 3C shows the average of N = 167 condensates recorded throughout the experimental volume, all with a radius of 1.98 µm. These averages reveal weak co-partitioning of phalloidin into the ICD condensates. Upon adding G-actin to the system, we observed diffusive mixing of G-actin throughout the sample. Although the presence of some degree of bleaching throughout the time course is apparent (see Fig. 3D and Supplementary Fig S5), the addition of G-actin to the solution was followed by a rapid increase in G-actin intensity inside the ICD condensates. After a characteristic lag time of approximately 18 min following the increase in G-actin signal, we observed an increase in phalloidin fluorescence inside the condensates, indicative of the onset of actin polymerization. Averaging images of condensates recorded throughout the experimental volume 90 min after addition of G-actin, we observed a pronounced ring of elevated phalloidin intensity at the interface of condensates, suggesting localization of emerging F-actin (Fig. 3E). This is in line with previous observations in polypeptide coacervates and protein condensates (*14*–*16*) and can be attributed to minimization of the bending energy of F-actin once the filament length exceeds the condensate diameter.

### Thermodynamic coupling of G-actin/F-actin inside PLPPR3 ICD condensates

The time-dependent phalloidin intensity inside condensates suggests two distinct regimes of F-actin polymerization kinetics (Fig. 3F). An initial regime of rapid polymerization is followed by a slower regime starting about 50 min after the addition of actin. The kinetics of the first regime are reminiscent of rapid polymerization kinetics observed in classical bulk solution assays (*17, 18*). However, while the F-actin concentration plateaus over long time periods in classical bulk polymerization, we observed a second slower, linear increase of F-actin inside PLPPR3 condensates. We attribute this behavior to dynamic coupling of G-actin partitioning from the surrounding liquid into the condensate and subsequent actin polymerization. Notably, in independent experiments, the second, linear polymerization regime is present, albeit not always as pronounced (Fig. 3H). In addition, photobleaching of G-actin and phalloidin during the time course (Supplementary Fig. S5) may lead to underestimation of actin enrichment.

G-actin is in thermodynamic equilibrium across the interface of ICD condensates if its chemical potential *μ*_*G*_ is equal inside and outside condensates. Thus, the ratio of equilibrium concentrations of G-actin inside and outside the condensate, *C*_*G,in*_ and *C*_*G,out*_, are fixed by the difference in standard chemical potential Δ*G*, which is the difference in affinity of actin to either phase:

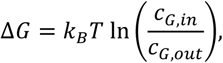

with Boltzmann constant *k*_*B*_, temperature *T* and assuming the chemical potential of an ideal gas.

Polymerization of G-actin into F-actin is a thermodynamic process that occurs when the critical concentration of G-actin, 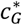, is exceeded. As F-actin forms, the concentration of G-actin inside the condensate decreases, leading to an imbalance in the partitioning of G-actin. This drives an influx of G-actin from the surrounding liquid to replenish G-actin now bound in filaments. This influx persists until the concentration of G-actin inside the condensate stabilizes at 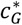, such that no further polymerization occurs. This will be the case if the concentration of G-actin outside the condensate is depleted to a value of

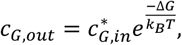

which satisfies the equilibrium condition for the partitioning of G-actin inside and outside the condensate. This model predicts an increase in overall actin intensity within the condensate over time and a decrease in actin concentration around the condensate, matching our experimental observations of increasing overall actin inside the condensate, as well as a continuous decrease in actin in the surrounding liquid at the onset of F-actin polymerization (Fig. 3F).

Polymerization kinetics of actin can be recapitulated using classical polymerization kinetics theory (*17*). Within this theoretical model, adding a small constant influx of G-actin to an existing pool of actin that exceeds 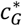 results in polymerization kinetics with similar regimes as observed in our data: an initial rapid polymerization regime (similar to that without added G-actin influx) is followed by a linear increase in F-actin (Fig. 3G). This model further suggests that the linear regime is characterized by elongation of existing filaments rather than nucleation of new filaments (Supplementary Fig. S6). In this way, PLPPR3 ICD condensates concentrate G-actin locally until conditions for F-actin polymerization are met, and subsequently sustain further polymerization by depleting G-actin from the environment.

Notably, in our model, a smaller degree of actin partitioning and/or higher 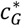 in the condensate do not inhibit the emergence of the secondary regime but slow the overall dynamics (Supplementary Fig. S6).

While large condensates lead to the formation of actin rings, we observe that small condensates, with a size similar to the resolution limit of our optical setup of about 200 nm, lead to outgrowth of straight filaments. F-actin filaments cannot comply with the strong curvature of such small condensates and will instead form straight, condensate-wetted bundles (Fig. 3I). The characteristic length scale at which filament localization changes is called the bendocapillary length (*19*).

### Molecular interactions between PLPPR3 and actin in condensates

According to our model, sustained F-actin polymerization inside PLPPR3 ICD condensates requires favorable partitioning of G-actin into the condensates, within which the G-actin concentration subsequently exceeds 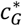. G-actin in the cytosol is in thermodynamic equilibrium with the actin cytoskeleton, i.e. the cytosolic G-actin concentration is at or close to 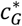. Assuming a similar 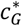 inside and outside of PLPPR3 ICD condensates, favorable partitioning of G-actin into PLPPR3 condensates would be sufficient to enable local F-actin polymerization. To investigate the molecular interactions between PLPPR3 ICD and actin inside and outside of condensates, we employed crosslinking mass spectrometry (XL-MS), a technique for analyzing protein structure and interactions (*20*). We treated mixtures of PLPPR3 ICD and G-actin, prepared with and without 5% PEG, with the lysine-specific crosslinker disuccinimidyl suberate (DSS). With a spacer length of approximately 12 Å and a maximum crosslinking distance of approximately 30 Å, DSS is able to link regions where the ICD and actin are in close proximity in bulk solution or under phase separation conditions, which are then revealed by subsequent mass spectrometry (Fig. 4A).

**Fig. 4.**
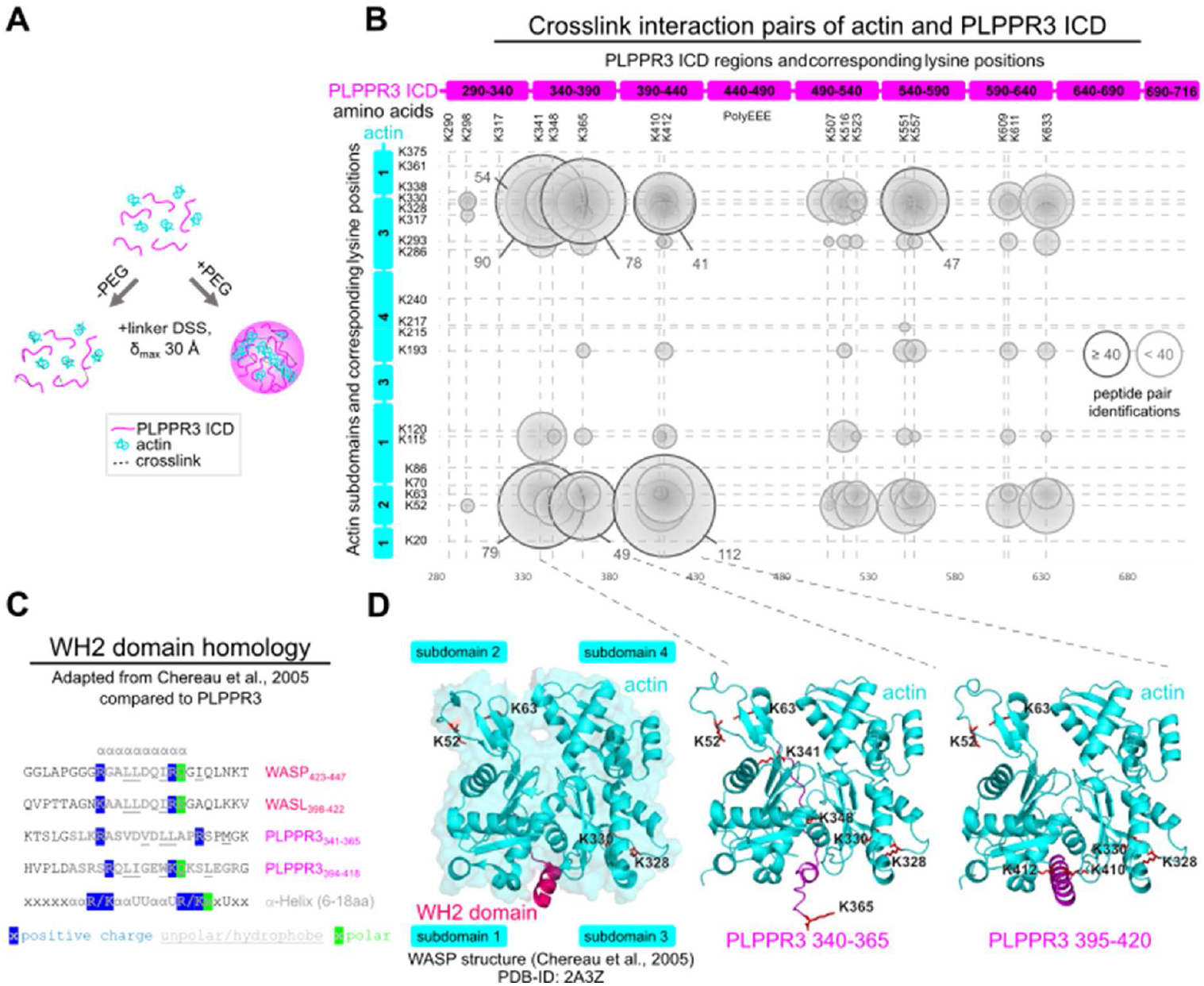
Crosslinking of condensates identifies possible PLPPR3-actin interaction domains. **(A)** Experimental design. Mixtures of PLPPR3 intracellular domain (ICD) and G-actin, in the absence and presence of 5% PEG, were incubated with the lysine crosslinker DSS with a spacer length of 12 Å. Crosslinking mass spectrometry (XL-MS) was used to analyze interacting regions. **(B)** Bubble diagram showing the number of identified crosslinked peptide pairs between PLPPR3 ICD and actin under PEG-induced condensate conditions (N=7), table format in Supplementary Fig. S10. **(C)** The highly crosslinked regions of PLPPR3 ICD (ICD 340-365 and ICD 395-420) have amino acid sequences similar to known WH2 domains (*21*). **(D)** AlphaFold3 prediction suggests that these PLPPR3 regions (magenta) are integrated into the hydrophobic ABC as a helix (PLPPR3 395-420, right) or a helix-like structure (PLPPR3 340-365, middle), similar to the WH2 domain of WASP (*21*). Note, that a broader amino acid sequence was used for modeling, while only the core WH2-domain homology was deleted for the ICD deletion variants ICDΔ345 and ICDΔ399 (see Materials and Methods).

Comparison of XL-MS data obtained with and without PEG revealed a significant increase in crosslinked peptide pairs between PLPPR3 ICD and actin upon PEG-induced phase separation. The most highly crosslinked regions of PLPPR3 ICD were amino acid (aa) residues 340-365 and 395-420 (Fig. 4B), suggesting that these regions may interact with actin. Sequence alignment of these PLPPR3 ICD regions revealed shared motifs with known WH2 domains, actin-binding domains found in the ABPs WASP and WASL (Fig. 4C). Both WASP and WASL also have an acidic region at some distance after the WH2 domain, similar to the poly-E-box in PLPPR3. Classical WH2 domains typically bind to the actin-binding cleft (ABC) between actin subdomains 1 and 3 (*21*). Using AlphaFold3, we modeled the highly crosslinked PLPPR3 ICD domains interacting with actin. These predictions suggest that the identified PLPPR3 ICD domains may integrate into the hydrophobic ABC as either helices (aa residues 395-420) or helix-like structures (aa residues 340-365) (Fig. 4D and Supplementary Fig. S8), closely matching WH2-actin interactions (*21*). Coherent with this, crosslinked PLPPR3 lysines were in close proximity to actin K328 and K330, located close to the ABC. Moreover, PLPPR3 lysines crosslinked to actin K52 and K63, which are in close proximity to the ABC in an actin filament. These results suggest that the ICD may contain WH2-like actin affinity domains.

Crosslinking experiments also support the findings from the actin polymerization assay. Crosslinks between actin lysines K115 and K193 are indicative of actin filament formation due to their respective distances in G-actin or F-actin. These crosslinks were not present in samples without PLPPR3 ICD. This was also true in the presence of PEG, suggesting only negligible levels of actin polymerization in the absence of PLPPR3 ICD (Supplementary Fig. S9).

### The actin affinity domain in PLPPR3 ICD affects actin partitioning and filopodia formation

To investigate the effect of the identified PLPPR3 WH2-like domains on actin partitioning, we purified PLPPR3 ICDs with deletion of amino acid residues 340-365 (hereafter ICDΔ345), 395-420 (hereafter ICDΔ399), or a double mutant (ICDΔ345/Δ399). All PLPPR3 ICD variants phase separated upon addition of 5% PEG, with partitioning coefficients comparable to wild-type ICD (Fig. 5A). We then quantified G-actin partitioning (*k* = *C*_*in*_/*C*_*out*_) into PLPPR3 ICD condensates, under latrunculin B treatment to inhibit actin polymerization.

**Fig. 5.**
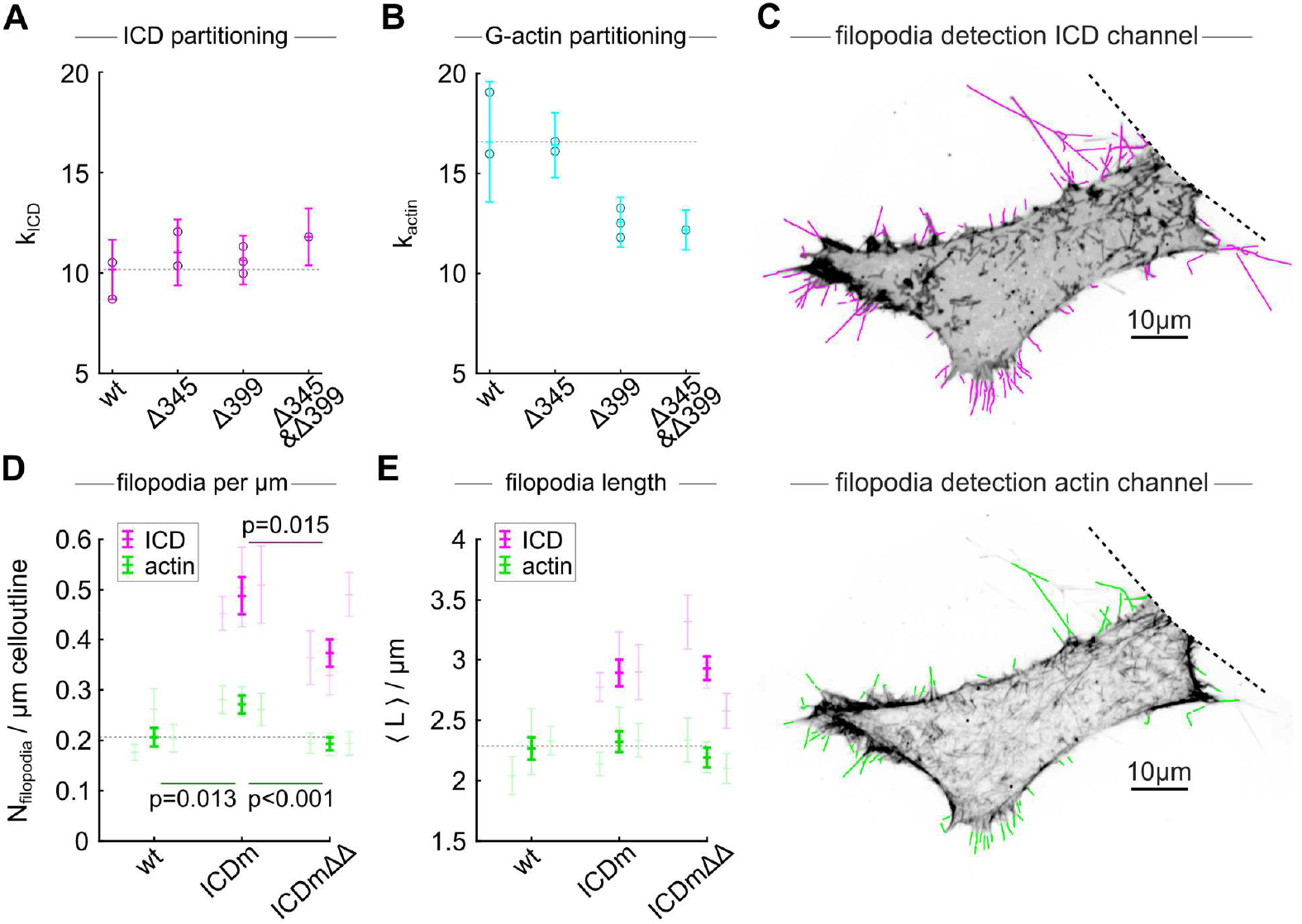
PLPPR3 enhances filopodia density in neuroblastoma cells. **(A)** Partitioning of wild-type ICD (N=4249 condensates from n=2 experiments), ICDΔ345 (N=3279; n=2), ICDΔ399 (N=3769; n=3) and ICDΔ345/Δ399 (N=846; n=1) to ICD condensates under 40 µM ICD and 1.2 µM G-actin treated with latrunculin B. Bars show overall mean ± SD, circles show means of individual measurements. Note that the standard error would be comparable to marker size due to the high N. **(B)** Partitioning of G-actin in the same experiments. The effect size between wild-type ICD and the ICDΔ399 variant in actin partitioning has a Cohen’s d of 1.71, indicating a strong effect. **(C)** Representative example of an N1E cell transfected with membrane tagged ICD (ICDm) analyzed by automated image analysis. **Top**: Image shows the PLPPR3 channel in inverted gray-scale. Filopodia registered in the image analysis routine are highlighted in magenta. **Bottom**: Corresponding filopodia detection in the actin channel. Filopodia registered in the automated image analysis routine are highlighted in green. The dashed lines indicate the cell segment cropped based on user input. **(D)** Average filopodia number density for N1E wild-type control, N1E transfected with ICDm, and N1E transfected with ICDmΔ345/Δ399. Data from filopodia detected via the actin and PLPPR3 channels are shown. Markers in the foreground show the overall mean and standard error, markers in the background show mean and standard error for each replicate. Mean and error were calculated considering the length of the cell outline in a given cell segment as the statistical weight with a total of N=72 segments in the actin channel of wild-type, N=48 for ICDm, and N=63 for ICDmΔ345/Δ399-transfected cells. P-values were calculated using two-sided Welch’s t-test considering the statistical weight. **(E)** Average filopodia length in the same datasets as for **D**.

While partitioning of G-actin into PLPPR3 ICDΔ345 condensates (*k*_*actin*_ = 15.2 ± 2.0, mean±SD) was unchanged compared to wild-type ICD (*k*_*actin*_ = 15.4 ± 3.0), partitioning of G-actin into PLPPR3 ICDΔ399 (*k*_*actin*_ = 11.8 ± 1.4) or PLPPR3 ICDΔ345Δ399 (*k*_*actin*_ = 11.8 ± 1.3) condensates was significant lower (Fig. 5B). These results indicate that the PLPPR3 region 395-420 mediates attractive interactions between PLPPR3 ICD and actin, possibly representing an actin affinity domain of physiological relevance. Within the framework of an ideal gas, the affinity of actin for the 399-domain is about 0.27 *k*_*B*_*T*, indicative of transient interactions. This interaction is expected to induce an up-concentration of G-actin in the condensate by a factor of 1.3 compared to the surrounding liquid. To probe the effect of PLPPR3 ICD on filopodia formation, we expressed membrane tagged (N-terminal myristoylated) PLPPR3 (ICDm) and mutant ICDmΔ345Δ399 variants in mouse neuroblastoma (N1E-115) cells. The cells were also transfected with eGFP as a cytosolic marker and transfection control and stained with SPY555-fastAct (Spyrochrome) to label actin.

Using an automated image analysis program (Fig. 5C), we observed that the number density of F-actin labelled filopodia increased from 0.21 ± 0.2 (mean±SE) filopodia per micrometer of cell outline in non-transfected cells to 0.27 ± 0.2 µm^-1^ in PLPPR3 ICDm-transfected cells (Fig. 5D). In cells transfected with the PLPPR3 ICDmΔ345/Δ399 mutant variant, the number density of F-actin labelled filopodia was comparable to non-transfected cells at 0.19 ± 0.02 µm^-1^. Accordingly, the number density of ICDmΔ345/Δ399-labelled filopodia was significantly lower compared to ICDm-labelled filopodia (Fig. 5D). In all cases, changes in number density of filopodia were not associated with significant differences in average filopodia lengths (Fig. 5E). The results analyzing the cytosolic eGFP staining as an alternative measurement are shown in Supplementary Fig. S11. These data demonstrate that PLPPR3 ICDm requires the WH2-like actin affinity domain in PLPPR3 to induce filopodia formation.

### Summary

Our work characterizes how a membrane protein can orchestrate filopodia formation via cytosolic liquid condensates. We show that thermodynamic coupling between G-actin partitioning into the condensates and subsequent localized actin polymerization sustain growth of F-actin inside condensates at the expense of G-actin in the environment. Because cellular F-actin networks share common G-actin sources (*22, 23*), F-actin inside condensates may thus outcompete cytosolic actin structures. Consequently, this mechanism allows the cell to form actin networks locally at the expense of pre-existing networks. This coupling of partitioning and polymerization inside condensates is functionally independent of filopodia regulating complexes via stochastic assembly of ABPs (*24*–*26*). The same mechanism may further contribute to previous observations of localized microtubule nucleation and growth in condensates of spindle assembly factors (*27, 28*). Overall, thermodynamic coupling of partitioning and polymerization of actin in condensates presents a novel mode of actin regulation.

While this mechanism offers an elegant handle for the cell to locally form actin networks *de-novo*, it may, in principle, be catastrophic for the integrity of the cytosolic F-actin cytoskeleton. Specifically, large condensates can retain substantial lengths of filaments by forming actin rings; however, condensates with radii below the bendocapillary length are unable to retain filaments. In our *in vitro* experiments, we observed this critical transition at condensate radii below approximately 200 nm (Fig. 3I). PLPPR3 condensates at the neuronal membrane are about half this size and are expected to have a lower surface tension compared to their *in vitro* counterparts (see coalescence experiments), further lowering their ability to retain bent filaments. These observations suggest that the small size of PLPPR3 condensates in cells is not incidental but reflects a self-regulatory mechanism that prevents filament retention and thus avoids potentially disruptive cytoskeletal reorganization in the cytosol.

We identified a WH2-like actin affinity domain within the PLPPR3 C-terminus and show that this domain is critical for induction of cellular filopodia, likely by mediating preferential partitioning. While the combination of favorable interactions between actin and PLPPR3 ICD and the ICD’s ability to form condensates alone is sufficient for PLPPR3 condensates to act as actin polymerization hubs, the function of PLPPR3 ICD condensates may be further boosted by favorable partitioning (or selective exclusion) of signaling molecules and/or ABPs (*4, 29*–*31*). In this way, membrane-bound cytosolic protein condensates present unique hubs of actin organization directly at and interacting with the plasma membrane.

## Acknowledgments

We thank Katrin Büttner, Kerstin Schlawe, Kristin Lehmann, Stephanie Bandura, Anja Koch, Brian Lally and Birgit Schroeer for technical assistance. Protein purification was supported by Laura Sandner. We thank Alexandra Polyzou for fruitful discussions. We acknowledge the support of the Advanced Medical BioImaging Core Facility (AMBIO) and the NeuroCure Multi-user Microscopy Core Facility of the Charité-Universitätsmedizin Berlin. Additionally, we thank Heike Stephanowitz from the Leibniz Institute of Molecular Pharmacology Berlin for XL-MS sample preparation, as well as Kathrin Textoris-Taube and Manuela Stäber for their advice on LC-MS sample preparation and analysis. We are grateful to Heike Nikolenko and Oxana Krylova from Leibniz Institute of Molecular Pharmacology Berlin for support with CD measurements. We thank Sarah Mackenzie for her professional assistance in editing and proofreading this manuscript. Parts of this work were published as a monograph as part of the requirements for obtaining a doctoral degree. The monograph is available from Free University Berlin, Germany at https://refubium.fu-berlin.de/handle/fub188/43424.

## Author contributions

Conceptualization: SB, TJB, WB, RLK, BJE

Methodology: TJB, SB, WB, BJE

Physical models and concepts: TJB

Protein purification: SB, MS, PS

Formal analysis: TJB, DNH, WB, SB

Experiments: SB, TJB, WB, DNH, PK, MR, VS, CK

Presentation: TJB, SB, WB, DNH

Funding acquisition: BJE, TJB, PS, RLK

Project administration: BJE, TJB, SB

Resources: BJE, PS, RLK, CMTS, FL, SW

Supervision: BJE

Writing – original draft: TJB, SB, WB, BJE

Writing – review & editing: TJB, BJE, SB, WB, GL

## Diversity, equity, ethics, and inclusion

We are committed to principles of diversity, equity, ethics and inclusion in all stages of our research processes and the communication of our findings. We strive to conduct and present research that respects the dignity, rights and contributions of all individuals, regardless of ethnicity, gender identity, sexual orientation, disability or cultural background.

## Data and materials availability

All data are available upon request and will be made available according to publication guidelines.

## Supplementary Materials and Methods

### Cloning and plasmids

The purification construct HA-M1-PLPPR3 ICD-His was gene synthesized (Eurogentec) including a 5’ Kozak sequence followed by a hemagglutinin signaling peptide (HA; A0M7P7) and a M1 Flag tag in front of the PLPPR3 ICD gene (amino acids residues 284-716). The 3’ end of the construct included a His_6_ tag, followed by a stop codon. The product was subcloned into a pCAX backbone (CAG promoter) using 5’ NheI and 3’ PstI restriction sites. Other DNA fragments were generated by a standard 35 cycle PCR amplification using KOD polymerase (#70086-3, Merck) and subcloned into the pCAX backbone using 5’-NheI and 3’-NotI as standard restriction sites. For PLPPR3 gene deletions, DNA fragments were designed with 10 bp overlaps that annealed in 10 cycles in a single tube extension (SOE) overlap PCR and were amplified by standard PCR (*32*). Each construct was validated by control digestion and sanger sequencing.

**Table S1.**
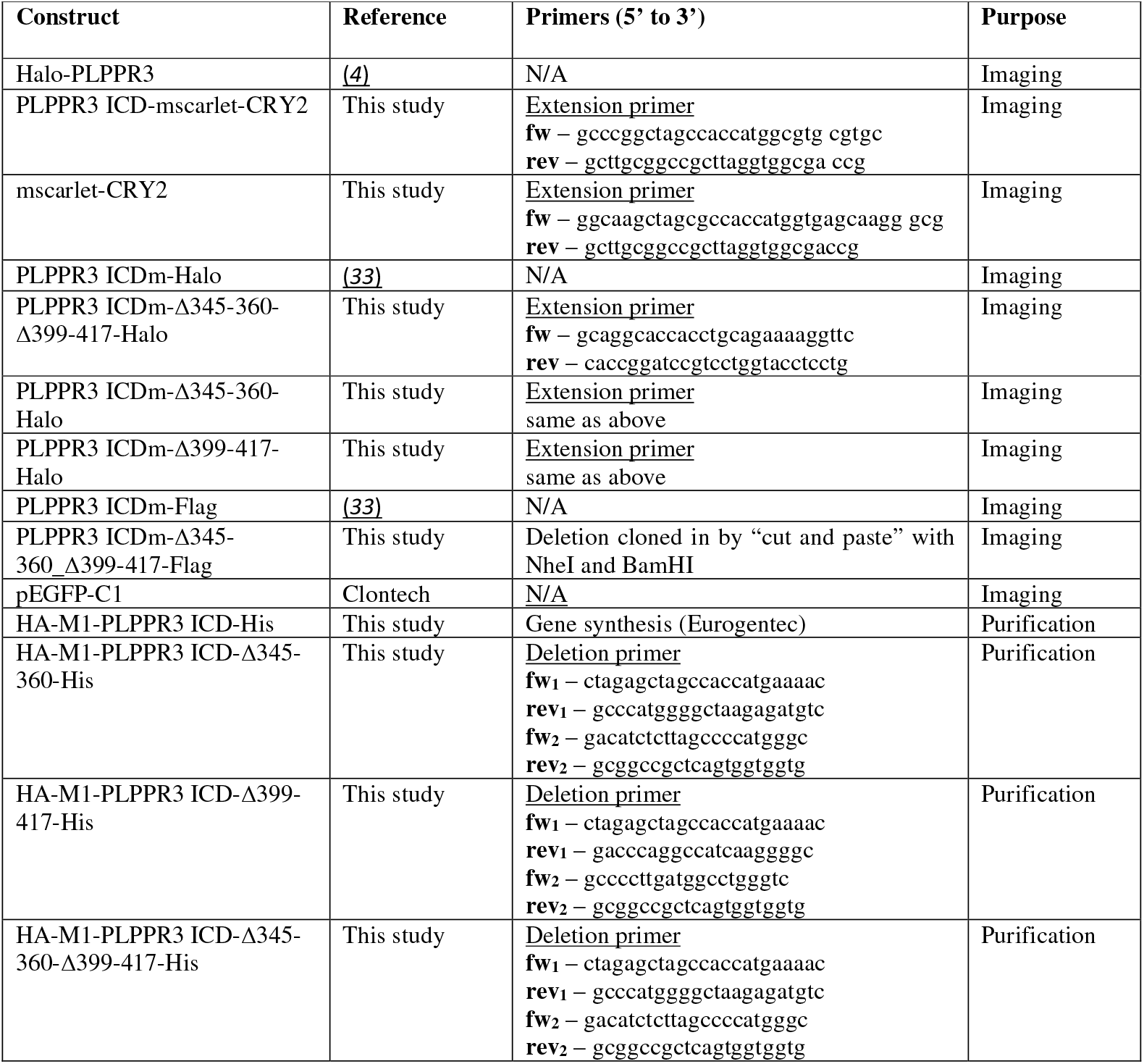
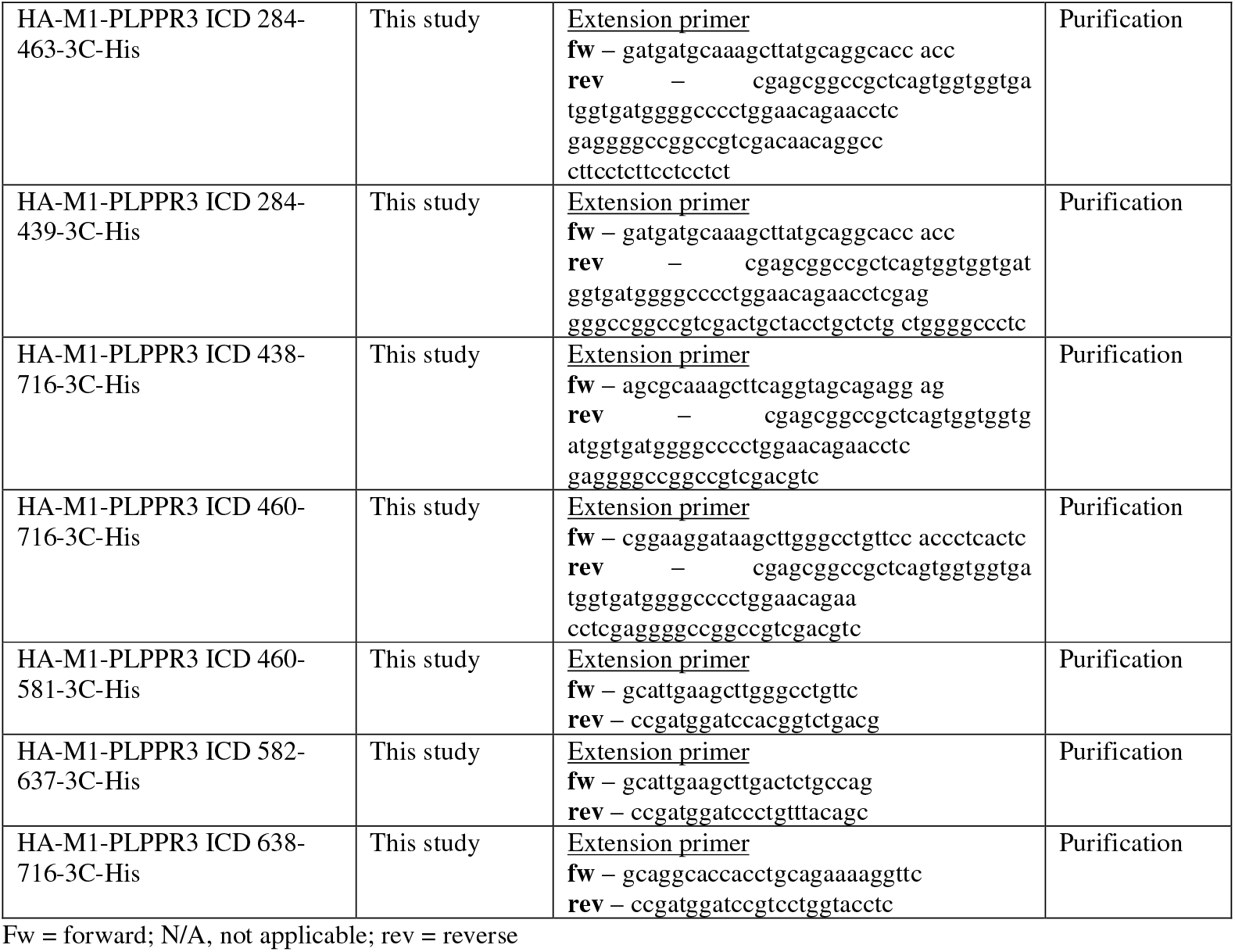
DNA constructs and constructing primers.

### Cell line culture and transfection

HEK293T (#CRL-3216) and N1E-115 (#CRL-2263) were purchased from ATCC and cultured in DMEM-high glucose (#31966047, Gibco) with 10% FBS (#P30-3031, PAN-Biotech) and 1% penicillin/streptomycin (#15140122, Gibco) at 5% CO_2_ and 37°C. Subculturing was performed twice a week until passage 25 with subcultures seeded at densities of 10,600 cells/cm^2^ and 13,300 cells/cm^2^, respectively. For protein purification the expression cell line Expi293F (#A14635, Thermo Fisher Scientific) was cultured in Expi293F Expression Medium (#A1435102) at 8% CO_2_, 37°C and 125 rpm. Subculturing was routinely performed twice a week up to 25 passages with subcultures seeded at densities of 0.5 to 0.6 x 10^6^ cells/mL.

For immunocytochemistry, 40,000 N1E-115 cells were plated on 12-well glass coverslips (Ø 18 mm) coated with 30 µg/mL poly-DL-ornithine (#P8638, Sigma) for 1 hour at 37°C, while for live imaging 80,000 HEK293T cells were plated on µ-slide 4-well glass bottom dishes (#80426, IBIDI) without substrate. Cells were incubated for 24 hours before transfection.

N1E-115 and HEK293T cells were transfected transiently with 1 mg/mL of Lipofectamine 2000 (#11668019, Thermo Fisher Scientific) in Opti-MEM reduced serum medium (#31985062, Thermo Fisher Scientific) with a lipofectamine to DNA ratio of 2:1 (v/v) per well. First, DNA and lipofectamine were separately incubated in 150 µL Opti-MEM for 5 min at room temperature.

Samples were mixed and incubated for 10 min at 37°C in a water bath. 300 µL transfection mix were added dropwise to the cell culture medium of each well. The transfection mix was incubated for 4 hours after which the cell culture medium was replaced with fresh medium. Cells were then grown for 24 hours before experiments.

### Protein expression and purification

Constructs designated for protein purification (Table S1) were transiently transfected into 2.5 x 10^6^ cells/mL Expi293F cell cultures with a PEI (1 mg/mL; #24765-100, PolySciences Inc.) to DNA ratio of 2.7:1 in OptiMem (#31985062, Thermo Fisher Scientific). The PEI:DNA complex was formed at room temperature for 25 min, before it was added dropwise to the culture. After 96 hours of overexpression, cells were pelleted at 4000x g for 5 min at room temperature and cell supernatant transferred to a fresh tube. The supernatant, containing overexpressed protein was flash frozen in N_2(l.)_ and stored at -80°C.

On the day of purification, supernatant was quickly defrosted in a water bath at 37°C and chilled on ice. For affinity chromatography, 2.5 mM CaCl_2_ (#2382, Merck) was added to the supernatant with 20 µL of M1 Flag sepharose bead slurry (in-house coupled) per mL of supernatant. Prior to this, beads were equilibrated with three washes of buffer containing 20 mM HEPES (#9105.3, Roth), 150 mM NaCl (#3957.7, Roth), 2.5 mM CaCl_2_ at pH 7.4 and sedimented at 500x g and 4°C. The supernatant was overhead rotated for 2 hours at 4°C with a speed of 14 rpm to allow beads to capture M1 flag tagged protein.

Loaded beads were separated from supernatant on a gravity flow column (#89896, Thermo Fisher Scientific), which was equilibrated with 20 mM HEPES, 150 mM NaCl, 2.5 mM CaCl_2_ at pH 7.4. Beads were washed three times with 5 column volumes (5 ml each) of a buffer containing 20 mM HEPES, 150 mM NaCl, 2.5 mM CaCl_2_ at pH 6.0 by gravity flow. For elution, 3 mL of elution buffer 20 mM HEPES, 150 mM NaCl, 0.2 mM Flag peptide (DYKDDDDK, GenScript), 5 mM EDTA (#X986.1, Roth) at pH 6.0 was incubated with beads for 30 min at 4°C. The eluted protein was captured in a 5 mL micro-centrifugal tube and 5 mM DTT (#04010.100, Biomol) was added. Proteins were filtered through PVDF membranes (#UFC30GV00, Merck) and concentrated (#UFC 803096, Merck) to 50 µL final volume. Protein purity was enhanced by size-exclusion chromatography using an Äkta Pure (GE Life Sciences) with a Superdex 200 increase 5/150 GL column (#28990949, Cytivia). The proteins were size separated in a buffer composed of 20 mM HEPES, 150 mM NaCl, 5 mM DTT at pH 6.0. Fractions containing high amounts of protein were combined and concentrated (#UFC503096, Merck) to 4-5 mg/mL, aliquoted in single use aliquots, flash frozen in N_2(l.)_ and stored at -80°C.

#### Protein fluorescence labeling

Freshly purified PLPPR3 ICD protein (500 µL at 1-2 mg/mL) was labelled fluorescently with DyLight^®^ 488 NHS Ester Dye (#46403, Thermo Fisher Scientific) in a buffer containing 20 mM HEPES and 150 mM NaCl at pH 6.0 for 1 hour at room temperature. Subsequently, excess dye was removed via dialysis (#71510-3, Merck) against a buffer containing 20 mM HEPES, 150 mM NaCl and 5 mM DTT at pH 6.0 overnight at 4°C. The labeled protein concentration and degree of labelling were determined using a nanodrop, aliquoted in single use aliquots, flash frozen in N_2(l.)_ and stored at -80°C.

### Primary hippocampal neuron culture and imaging

All animal procedures were carried out under the license T0347/11 in accordance with the regional health authority. Primary hippocampal neurons were dissected (sex unspecific) on embryonic day E16.5 from wildtype mice with a C57 BL/6NCrl background. Hippocampi were isolated and briefly pre-incubated in micro-centrifugal tubes (two hippocampi/tube) containing cold HBSS (#14170-070, Thermo Fisher Scientific). Tissue was rinsed with fresh HBSS, then digested with 500 µL Papain (20 U/mL; #LS003126, Worthington Biochemical) for 30 min at 37°C, shaking gently at 350 rpm. The reaction was quenched by prewarmed stop solution composed of DMEM with 10% FBS, 1% penicillin/streptomycin, 2.5 mg/mL BSA (#A2153, Sigma) and 2.5 mL/mg trypsin inhibitor (#T9253, Sigma) for 5 min at 37°C. The medium was aspirated and replaced with 200 µL plating medium/tube composed of Neurobasal Media (#21103-049, Thermo Fisher Scientific), 10% heat inactivated FBS, 2% B27 (#17504044, Thermo Fisher Scientific), 1% GlutaMax (#35050-038, Thermo Fisher Scientific) and 1% penicillin/streptomycin. The tissue was gently disrupted by trituration and neurons of different preparations pooled into a single tube. After counting, neurons designated for immunocytochemistry experiments, were plated in plating medium on 12-well glass coverslips (Ø 18 mm) coated with poly-DL-ornithine (30 µg/mL) and laminin (20 µg/mL; #L2020, Sigma), with a density of 120,000 neurons/well. After 2 hours the plating medium was replaced by complete medium composed of Neurobasal Media, 2% B27, 0.5% GlutaMax and 1% penicillin/ streptomycin.

Primary hippocampal neurons were transfected at day *in vitro* 1 (DIV1) using Ca^2+^ phosphate transfection (*34*). In brief, 500 µL/well Neurobasal Medium containing 2% B27 and 0.25% GlutaMax were prepared and pre-warmed to 37°C. The transfection solution was formulated by diluting a 2 M CaCl_2_ stock (2.5 µL/well) with ddH_2_O (22.5 µL/well), then adding DNA (4 µg/well; 2 µg each/well for co-transfection) and mixing thoroughly. A 2x BSS Buffer (25 µL/well; 50 mM BES, 280 mM NaCl, 1.5 mM Na2HPO4, 2 mM pH 7.26) was added drop by drop and the solution gently mixed by inverting. Finally, 450 µL/well of the pre-warmed Neurobasal Medium were added and the transfection mix incubated 15 min at 37°C in a water bath. The neuronal medium was collected, kept tempered at 37°C and exchanged with the transfection mix for 20 min at 37°C. The transfection mix was aspirated, each well washed three times with 500 µL pre-warmed HBSS buffer (135 mM NaCl, 4 mM KCl, 1 mM Na_2_HPO_4_, 2 mM CaCl_2_, 1 mM MgCl_2_, 20 mM HEPES, 20 mM D-glucose, pH 7.3) and the neuronal medium added back into each well. The primary cultures were incubated at 37°C and 5% CO_2_ until the indicated DIV.

#### Immunocytochemistry

After 24 hours post-transfection, cells or neurons were washed once with PBS (#BE0014G, Oxoid) and fixed with pre-warmed 4% paraformaldehyde (#104005100, Merck) for 15 min at room temperature. PFA was aspirated and cells washed three times for 5 min with PBS. Neurons were antigen-retrieved by adding 800 µL/well of citrate buffer (10 mM citrate acid #A7496.0500 Applichem, 0.05% Tween-20 #655205, Calbiochem at pH 6.0), incubated for 15 min at 92°C and washed once with room temperature PBS. Fixed cells or neurons were permeabilized with 0.1% Tween-20 for 10 min at room temperature and non-specific binding blocked with 4% goat serum (#16210-072, Thermo Fisher Scientific) in PBS for 1 hour at room temperature. The primary antibodies (Table S2) were diluted in blocking buffer and incubated overnight at 4°C. The following day, cells or neurons were washed three times for 10 min with PBS. The secondary antibodies (Table S2) were diluted in blocking buffer and incubated for 1 hour at room temperature. The cells or neurons were washed two times with PBS and the third time with PBS plus Hoechst stain (Table S2). The cells were mounted with Prolong Glass Antifade Mountant (#P36984, Invitrogen) on imaging slides (#J1800AMNZ, Thermo Fisher Scientific) and stored at 4°C.

#### Imaging of endogenous PLPPR3 in fixed neurons

DIV5 hippocampal neurons were either untreated, treated or treated/treatment washed-off prior to fixation and permeabilization. For treatment, neurons were incubated with 3.5% (v/v) 1,6-hexanediol (#240117, Sigma) for 5 min at 37°C. The treatment was washed off with two washes of PBS, before the medium was changed into Neurobasal Medium (low GlutaMax) for 2 hours at 37°C. Following, PLPPR3 was probed by α-PLPPR3 (1:5000) together with the axon marker α-Tau (1:5000) and visualized by secondary α-rabbit 488 (1:500) and α-chicken Cy5 (1:500), respectively. HOECHST dye (Table S2) was used to visualize nuclei. Imaging was performed on a Nikon A1Rsi+ confocal system (Nikon) with a 60x oil-immersion objective (NA 1.4). Single axons were chosen based on the Tau staining and imaged with 405 nm, 488 nm and 647 nm laser channels at 0.7% (0.0308 mW), 5% (0.04 mW) and 1.6% (0.272 mW) laser intensity, respectively. Images were processed with Fiji (Image J v1.53t).

#### Live visualization of filopodia in neurons

Primary hippocampal neurons transfected with 0.3 µg UTR-GFP and 0.5 µg Halo-PLPPR3 were imaged as described in (*4*). In brief, neurons were labelled for 15 min with Janelia Fluor 646 HaloTag ligand (Promega) at a final concentration of 100 nM. The medium was replaced with Neurobasal A minus phenol red (#12349015, Gibco), supplemented with 2% B27, 1% penicillin/streptomycin, 1% GlutaMax and 100 µ M β-mercaptoethanol (#M7522; Sigma). Human recombinant BDNF (50 ng/mL, #248-BD, R&D systems) was added to the medium to enhance filopodia growth. Cells expressing a medium amount of the constructs were selected for imaging with a spinning disk confocal CSU-X (Nikon) equipped with a 100x oil-immersion 1.45 NA objective under controlled 37°C, 5% CO_2_ and humid environment. Axons were imaged for 10 min (8 s/frame, 75 frames) with 488 nm and 647 nm laser lines at 300 ms exposure scanning in z-direction.

### Antibodies and dyes

**Table S2.** Antibodies and dyes.

| Antibody/Dye | Manufacturer | Catalog # | Species | Dilution | Application |
| --- | --- | --- | --- | --- | --- |
| PLPPR3<br>1.54 μg/μL | Custom-made<br>(Eurogentec) | - | rabbit | 1:5000<br>1:1000 | ICC<br>WB |
| α-Tau | Abcam | #ab75714 | chicken | 1:5000 | ICC |
| α-MAP2 | Abcam | #ab5392 | chicken | 1:5000 | ICC |
| Donkey-α-chicken<br>Cy3 | Dianova | #703-165-155 | donkey | 1:500 | ICC |
| Donkey-α-chicken<br>647 | Dianova | #703-605-155 | donkey | 1:250 | ICC |
| Phalloidin 488 | Sigma | #U9409 | N/A | 1:1000 | ICC |
| Phalloidin 565 | Merck | #94072 | N/A | 1:1000 | IVPS |
| Janelia Fluor 646<br>HaloTag ligand | Promega | #GA1120 | N/A | 1:1000 | LCI |
| Janelia Fluor 549<br>HaloTag ligand | Promega | #GA1111 | N/A | 1:1000 | LCI |
| Hoechst | Sigma | #14530 | N/A | 1:20,000 | ICC |
| α-rabbit IgG HRP | Vector Laboratories | #AB_2916034 | goat | 1:5000 | WB |
| α-rabbit-488 | Jackson<br>ImmunoResearch | #AB_2307443 | donkey | 1:500 | ICC |
| Spy555-FastAct | Spirochrome | #CY-SC205 | N/A | 1:1000 | LCI |
| Verapamil | Spirochrome | #CY-SC012 | N/A | 1:1000 | LCI |

ICC – immunocytochemistry; WB – western blot; IVPS – *in vitro* phase separation; LCI – live cell imaging; N/A – not applicable

### Live cell imaging and analysis of PLPPR3 in HEK

#### CRY2 fluorescence recovery after photobleaching (FRAP)

HEK293T cells were transfected with PLPPR3 ICD-mscarlet-CRY2 and imaged with a spinning disk confocal CSU-W1 SoRa (Nikon) equipped with a Plan Apo 60x NA 1.4 oil immersion objective (Nikon) and a heating chamber set to 37°C and 5% CO_2_. CRY2 oligomerization was activated with 1% 488 nm laser (0.103 mW) for either 1 s or 5 s. Immediately after this activation, time lapse imaging began, lasting for 5 min (1 s frame rate; 30 frames). Complete condensates were bleached using the 555 nm laser and subsequently imaged at one frame per second.

FRAP was analyzed via ImageJ by recording the intensity in the region of interest compared to a reference in the cytosol.

#### CRY2 coalescence

Transfected HEK293T were also used for coalescence experiments. Images were taken on the same microscope at 0.02 s between frames using the 4x SoRa mode with the 60x oil immersion objective. Coalescence events were detected by visual inspection of the data and cropped for automated analysis in Matlab using a custom image analysis pipeline. Data are upscaled twofold using bilinear interpolation to minimize errors due to small cluster size and blurred with a Gaussian filter to decrease shot noise. PLPPR3 condensates were registered if the local intensity exceeded an initial heuristic threshold value of four standard deviations above background level, which was identified via the median intensity for each frame. Registered pixel groups were thus characterized using the regionprops function. An intensity profile across each condensate’s interface was calculated based on each pixel’s minimal distance to any pixel within the detected condensate’s interface (see also (*8*)). This intensity profile allows determination of a corrected threshold value as the intensity at the maximal gradient in intensity across the interface. The initial thresholding value was chosen to closely match the correct value to ensure that condensates are correctly registered. The principal axis ratio of the major to minor principal axis of the detected blob was recorded over time. The onset of coalescence was registered as the timepoint at which the principal axis ratio increased and remained elevated in the consecutive frame. Images show original data; overlain outlines are based on the upscaled analysis (see Fig. 1D and 2C).

We employed a simple description of coalescence dynamics based on dimensional analysis. Liquid droplets coalesce due to minimization of the surface energy. This process is faster, the higher the surface tension *γ*, whereas the resulting flow is damped by the liquid’s viscosity *η*. The ratio *η*/*γ* is known as the inverse capillary velocity and characterizes the relaxation dynamics. This inverse velocity can be approximated via the time scale of relaxation of the shape *τ*, obtained from fitting an exponential to the PA ratio over time, and a characteristic length scale of the flow, corresponding here to the radius *R* of a given condensate, estimated as 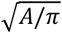, with the cross-sectional area of the condensate after coalescence *A*. This consideration yields *η*/*γ* ≈ *τ*/*R*. For a more detailed analysis refer to (*35*). Note that the viscosity of the surrounding medium also effects the relaxation dynamics. In fact, some coalescence events show two timescales of relaxation, one fast and well approximated by an exponential, one slow and almost linear. This hints at initial droplet coalescence and subsequent (viscoelastic) relaxation of the surrounding cytoplasm. To limit the impact of the secondary regime on the coalescence dynamics, we only considered the first 2 s after the onset of coalescence in the exponential fit used to extract *τ*.

### PLPPR3 ICDm and ICDmΔ345/Δ399 transfection and imaging in N1E cells

N1E-115 cells were transfected with either eGFP; eGFP and PLPPR3 ICDm-Halo; or eGFP and PLPPR3 ICDmΔ345Δ399-Halo. Janelia Fluor 646 HaloTag Ligand (Promega, #GA1120) was added to the cells at a final concentration of 200 nM, 1.5 hours before imaging and incubated for 30 min. The medium was replaced with fresh complete medium containing Spy555-FastAct (Spirochrome, #CY-SC205) and Verapamil (Spirochrome, #CY-SC012) following the manufacturer’s protocol (dilution 1:1000 in medium). Following a 1-hour incubation, cells were imaged in this medium. Live cell imaging of confocal stacks with a step size of 0.3 µm was done on a Nikon SoRa Spinning Disk Confocal CSU-W1 microscope fitted with an Okolabs incubation chamber set to 37°C with 5% CO_2_ during imaging. Cells were imaged in the z-direction from bottom to top using 488 nm (eGFP) and 638 nm (Halo dye) laser lines with 10% (1.03 mW) and 10% (0.82 mW) of power respectively with 200 ms exposure for both lines.

#### Filopodia analysis

Analysis of filopodia was carried out in a custom-made Matlab analysis pipeline. The location of the interface between cell and coverslip (cell base) was determined by calculating the median intensity for each z-slice of the confocal stack plus the 80^th^ percentile in each slice. The position of the maximal gradient in this metric reliably detected the cell base. Confocal stacks were transferred to maximum projections considering four z-slices from the cell base upward, corresponding to a height of 1.2 µm.

Segments analyzed were selected by user input to identify single cells and avoid staining artifacts or defects. For each input, the cell body was determined through the cytosolic eGFP stain. Thresholding was done automatically for each segment based on the intensity histogram. The intensity histogram was calculated with 200 bins between the minimum in each image and the 95^th^ percentile. The resulting histograms were smoothed to decrease noise. The threshold was determined as the first local minimum after the first maximum for each channel. Because the first maximum represents the background, this metric reliably separated the cell from background. The resulting mask of the cell body was processed using structural elements to close small holes and smooth edges. This step primarily served to determine the rough mask of the cell. The intensity of eGFP, ICD and actin inside and outside the cell were determined as the trim-mean (trimming 80% of data) of pixels inside and outside the mask applied to the respective channel. Cells with an expression level below four in the eGFP channel were discarded due to uncertainty in the cell body detection. Measurements for a given channel were only considered if the respective ratio of intensity inside to outside the cell also exceeded a value of four.

Filopodia in each channel were detected via a difference of Gaussians of the respective channel. Filopodia were registered if the intensity in this metric exceeded 1/3 of the standard deviation in the respective image. This method is sensitive to gradients but not to extended structures, the resulting mask was thus combined with the cell mask to also accurately detect the volume of the cell. Registered pixel clusters containing less than 50 pixels were discarded.

The resulting map of registered pixels was further filtered for accurate detection. First, all groups of registered pixels that did not connect to the cell body mask dilated by a disc with 14 pixels radius (1 µm) were discarded. This ensured that all registered filopodia originate from the cell in question. Second, the resulting map of connected structures was subsequently inflated by a disc of 7 pixels radius (0.5 µm). Registered groups of pixels in the initial map that overlapped with this inflated mask were also accepted. This method patched small dark segments along a filopodium as well as crossing filopodia that were typically interrupted due to the difference of Gaussian detection. This final mask of accepted pixels was used to determine filopodia as pixels that disappear upon morphologically opening the mask with a disk of 5 pixels radius. 10 pixels were the upper diameter of structures considered as filopodia based on optical inspection of the data.

The final filopodia were skeletonized. The number of filopodia was determined as the number of end points of skeletonized filopodia that originated from within 1 µm of the cell body minus the number of these filopodia. This ensures that a filopodium with a break due to a dark spot is not counted twice. Moreover, this ensures that filopodia that are connected, e.g. because they originate from the same position on the cell outline (within our optical resolution) or because they branch, are each considered (e.g. two filopodia emerging from the same base are counted as two).

The average length of filopodia was calculated as the total number of pixels in all skeletonized filopodia divided by the number of filopodia (see above) for each cell segment. Note that this method neglects the fact that diagonal pixels have a distance of 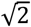, whereas direct neighbors have a distance of one. This method consequently tends to underestimate filopodia length, but is well suited to compare conditions.

All presented averaged values have been weighted by the cell outline of the respective cell segment to ensure equal weight per unit cell outline. In this way, analyzing one cell as a whole, or separating it into multiple segments, contributes the same to the final average value.

For the actin channel, a total length of cell outline of 1.1 cm has been analyzed for control cells, a total outline of 0.8 cm for ICDm-transfected cells and of 1 cm for ICDmΔ345Δ399-transfected cells.

### Condensate imaging in vitro

Round imaging dishes (#6160-168, Miltenyi Biotec) were prepared by plasma-cleaning the surface (5 min at 40% power) and coating with 1% (w/v) PVA (#363065, Sigma) for 20 min at room temperature. PVA solution was removed, and dishes dried overnight at an upright angle of 45°. Before imaging sessions, dishes were briefly cleaned with compressed air and a moist kimtech wipe (#830-34155, MG chemicals) was draped around the imaging surface to ensure a humid environment.

Protein condensates were generally formed in 0.5 mL micro-centrifugal tubes (#0030108035, Eppendorf) with PLPPR3 ICD (including 3% labelled PLPPR3 ICD using NHS-ester conjugated to Alexa-488) in protein buffer (20 mM HEPES, 150 mM NaCl, 5 mM DTT at pH 6.0), 1:10 (v/v) of 10x F-actin buffer (10x is 1 M KCl, 0.1 M imidazole, 10 mM ATP, 20 mM MgCl_2_ at pH 7.4), and 5% (w/v) PEG8000 (#V3011, Promega) for a total volume of 4 µL. Only the phase diagram was acquired without F-actin buffer and at varied PEG concentrations. Note that phase diagram experiments were done at specified concentrations in mg/mL and thus do not align with ticks in molar concentration.

Imaging was performed on a spinning disk confocal CSU-W1 SoRa (Nikon) equipped with a Plan Apo 60x NA 1.4 oil immersion objective (Nikon) and perfect-focus system (PFS) switched on for timelapse experiments. In all setups, images of the microscope background noise were taken before or after the experiment.

#### In vitro FRAP

Condensates were formed at 20 µM PLPPR3 ICD in the presence of protein buffer, F-actin buffer and 5% PEG (w/v). A 488 nm laser line was used to photo bleach a fraction of given condensates (partial FRAP). The imaging loop consisted of a pre-bleach baseline (5 s frame rate, 3 frames) at 2.5% 488 nm laser power (0.257 mW), a bleach phase for 2 s at 70% 488 nm laser power (7.2 mW) and post-bleach timelapse (5 s frame rate, 120-240 frames) at 2.5% 488 nm laser power. Several condensates in a field of view were left unbleached and used as references to account for subsequent bleaching throughout the post-bleach timelapse. Moreover, background images were taken on the imaging dish but outside the experiment volume to correct for the noise floor of the imaging setup.

Data were analyzed using custom Matlab pipelines. Photobleached condensates were detected by the difference between the frame before and after photobleaching. Reference (unbleached) condensates were automatically detected as those condensates that were not bleached and that stayed in one location throughout the image acquisition, i.e. had pixels that stayed within the initial condensate mask at all times. Tracking of condensates migrating short distances was implemented to follow reference condensates. If no condensates were sufficiently stationary, a user input prompt asked for manual definition of reference condensates that were than tracked automatically. Following an initial rough thresholding based on the intensity values throughout the data, a radial intensity profile from the center of each condensate was calculated. These profiles were used to define a mask to extract the intensity inside each condensate, considering only pixels within a radius at which the radial intensity was above 90% of the intensity in the center. Intensities for reference condensates were then calculated as the mean over the intensities within this mask. A binary mask of each condensate at timepoint *t* was dilated and used as a mask for condensate detection at timepoint *t+1* to allow for some degree of motion of the condensates. The shift in position for each condensate was recorded over time to check for consistency. The mean intensity of reference condensates typically showed weak signs of photobleaching. In order to smooth the reference intensity profile taken as the mean across reference condensates, the reference profile was fitted with a second order polynomial.

In photobleached condensates, additionally the bleached region was registered as the region not exceeding 10% of the minimal intensity immediately after photobleaching. This mask of the initial bleach spot was imposed on all subsequent timepoints to evaluate the intensity at the bleaching site. Immediately after photobleaching, the automated detection of condensates was initially prone to give faulty results if the bleached spot was close to the edge. This effect was mitigated by thresholding subsequent frames on the assumption that the condensate size remains constant. This was done by using a percentile intensity cutoff to ensure that the same number of pixels remains in the condensate at all times.

In addition, the maximal intensity throughout the entire bleached condensate was recorded. Rapid increases in this metric indicated coalescence of other condensates into the photobleached condensate. Data for the corresponding condensate were then cut at the timepoint before coalescence.

The intensity profile within the bleached spot for each such condensate was divided by the reference profile to account for subsequent photobleaching throughout imaging and re-scaled to a value of one, compared to pre-FRAP. This profile was then fitted with the sum of two exponentials to extract fast and slow recovery dynamics.

Accurately extracting the diffusion coefficients from such data is challenging due to the geometry and spatial dimension of the problem, as well as largely unknown dynamics across the interface and thus uncertainty in the boundary conditions. Here, we used a minimal model to extract only apparent diffusion constants that allow for quantitative comparison between measurements and a rough order of magnitude estimate of the actual diffusion constants. For this we utilized the relation *τ*∼*R*^2^/*D*, with diffusion time *τ*, diffused distance *R*, and diffusion constant *D* (*36*).

The internal diffusion coefficient *D*_*app,in*_ was then calculated using time *τ* from the fast exponential fit and *R* as the radius of the initially bleached spot. The external diffusion constant *D*_*app,out*_ was then calculated using time *τ* from the slow exponential fit and *R* as the radius of entire condensate.

While we employed strong simplifications in this quantification, the result *D*_*app,in*_ ≈ 0.05 *μ*m/s^2^ appears reasonable. An internal diffusion constant of 0.05 µm^2^/s was also reported for the silicone oil poly(dimethylsiloxane) (PDMS) consisting of 400 dimethylsiloxane molecules, each with a molecular weight of 74 Da, at 25°C (*11*). The ICD consists of 430 amino acid residues with an average molecular weight of 110 Da and thus shows a roughly similar molecular architecture. Note, however, that the protein condensate is solvated by water, whereas the oil is not.

#### In vitro coalescence

Condensates were formed at 20 µ M PLPPR3 ICD (including 3% PLPPR3 ICD labelled with Alexa 488) in the presence of protein buffer, F-actin buffer and 5% (w/v) PEG for a final volume of 4 µL. First experiments were done without ATP, later experiments with 1 mM ATP. The timepoint at which PEG was added was recorded for each experiment as a reference point. Time-lapse images were acquired at a frame rate of 20 ms at 5% (0.514 mW) laser power in the 488 nm line with a duration of 5 min using a Plan Apo 60x NA 1.4 oil immersion objective (Nikon), 2x2 binning and fast scan mode. The field of view was changed between timelapse experiments. Coalescence events were cropped out of the original data following visual inspection and analyzed similar to CRY2 coalescence experiments *in vivo*. Condensates were initially registered by a heuristic threshold. Based on this initial registration, a radial intensity profile was calculated for each condensate before the onset of coalescence and the corrected threshold was set to the value corresponding to the maximal gradient in intensity across the condensate interface. Subsequent analysis recorded the principal axis ratio throughout the coalescence, fitted an exponential to extract the characteristic time scale and utilized the cross-sectional area of the final condensate to estimate the characteristic length scale, as for PLPPR3 ICD-CRY2 coalescence analysis (see above). Data were considered outliers if the final aspect ratio given by the offset of the exponential fit was below 1 (unphysical) or above 1.05 (suggesting imperfect relaxation, e.g. due to adherent lint or wetting to defects in the surface coating). Data with a fit that yielded a coefficient of determination below 0.97 were also discarded, as were outliers in relaxation time, defined by being outside 1.5 standard deviations of the mean. Out of 134 events, these criteria identified 14 outliers.

#### Actin partitioning assay

Experiments were carried out with 40 µM PLPPR3 ICD (including 3% PLPPR3 ICD labelled with Alexa 488) and deletion variants in protein buffer, F-actin buffer, Latrunculin B (#3974, Tocris, final concentration 5 and 10 µ M), 1.2 µ M actin and 5% PEG in a total volume of 4 µL. An elevated concentration of ICD was used compared to other experiments to achieve large condensates. The composition was thoroughly mixed and a 2.5 µL droplet pipetted onto an imaging dish coated with PVA. At random fields of view, entire condensates were scanned in z-direction from bottom to top using 488 nm (PLPPR3 ICD) and 638 nm (actin) laser lines with 2.5% (0.257 mW) and 5% (0.41 mW) of power respectively. Imaging was carried out as long as 2 hours after addition of PEG and revealed consistent partitioning over time. Background images were taken outside the experimental volume.

A custom Matlab analysis pipeline was used to analyze the partitioning. First, all data were corrected by the background from the optical instrument. Condensates were then registered via thresholding within the confocal stacks. Initially a rough threshold value was determined based on the mean intensity and standard deviation in the 488 nm channel for the ICD. Due to the presence of small drift in some samples, the image in the actin channel was detected in a box surrounding the ICD channel to allow for drift up to 10 pixel (1.1 µm) during subsequent imaging of the different channels. The global threshold for the actin channel was determined such that the total number of registered voxel (i.e. total condensate volume) matched the total volume in the 488 channel. The difference in volume and location for each individual condensate was recorded and outliers discarded to ensure consistent matching of condensates in each channel. For each channel, an image of the xy-plane closest to the centroid of a given condensate and extending at least 20 pixels from the condensate interface was extracted and analyzed further. In these images, the surface-relative intensity profile was calculated for each channel as for coalescence experiments, see above and (*8*). Based on this metric, the accurate interface position was determined as the location with maximal gradient across the interface. The interface width was determined as the region across the interface, in which the local intensity gradient after smoothing exceeded 15% of the maximal gradient. The interface width was then taken as half of this value. The intensity outside (inside) a condensate was then taken as the mean of all pixels outside the interface position minus (plus) the interface width. The ratio of intensity inside to outside determined the partitioning for a given condensate in a given channel. Other values that were extracted and recorded were the condensate volume and the time after addition of PEG for each dataset. Condensates with a radius below 1.5 µm were not considered because smaller condensates showed a dependence of the partitioning value on condensate size, likely due to optical artifacts. Partitioning in condensates above 1.5 µm radius was independent of size for actin and only weakly dependent on condensate size for ICD.

#### Actin polymerization assay

Condensates were generated at 20 µ M PLPPR3 ICD (including 3% PLPPR3 ICD labelled with Alexa 488) with protein buffer, F-actin buffer, phalloidin 565 (final concentration 20 nM unless stated otherwise), 5% (w/v) PEG, and 1.2 µ M alpha-actin (#8101-01, Hypermol) including 31% atto-647 actin (#8158-01, Hypermol), in a total volume of 4 µL. All components except actin were combined in a micro-centrifugal tube, pipetted on the imaging dish and incubated for at least one hour for condensates to sediment. Before addition of actin, several confocal stacks of condensates were taken, as well as images of the background outside the experimental volume. Actin was gently pipetted into the middle of the droplet formed by the sample and timelapse imaging started immediately. For the setup, the laser lines 488 nm (PLPPR3 ICD), 561 nm (phalloidin) and 638 nm (actin) were used at 2.5% (0.257 mW), 5% (0.53 mW) and 5% (0.41 mW) respectively. The framerate was set to 1/30 s while scanning through the condensates in z-direction at 0.4 µm/step (10 µm total). After the time lapse, confocal stacks of condensates outside the field imaged during the timelapse were taken. For experiments longer than 90 min, the framerate was reduced to 1/300 s after the first 30 min to reduce photobleaching.

Analysis was carried out in a custom-made analysis pipeline in Matlab.

Confocal stacks taken before and after the timelapse, as well as each confocal stack within the timelapse, were analyzed in the same way as data for partitioning (see above), with addition of the 561 nm channel for phalloidin.

Throughout the timelapse and in the post-timelapse, some condensates, typically small, showed very bright phalloidin and actin signals. Inspection of the timelapse revealed that these condensates fell from higher up in the experimental volume, already bright in actin and phalloidin. We suspected that these condensates were closer to the point of actin injection and were subsequently subject to high actin concentrations. We suspect that the mechanism for actin polymerization at the expense of G-actin in the environment proposed in the main text is responsible for creating these highly actin-rich condensates. Since the equilibrium partitioning conditions only define a ratio of G-actin inside and outside the condensate, a very high local G-actin concentration (as expected close to the injection point) will lead to an even higher concentration inside condensates, giving rise to very pronounced polymerization in such condensates. To minimize the number of free-floating condensates, condensates were allowed to settle for at least one hour before addition of actin. The remaining bright actin condensates were detected using the Matlab function *isoutlier* based on the median of the phalloidin intensity throughout all condensates, as well as, the standard deviation of the phalloidin intensity.

Note that a comparison of the intensity inside condensates at the end of the timelapse compared to the intensity throughout confocal stacks after the timelapse revealed bleaching also in the actin and phalloidin channels (see Fig. 3D, E and Supplementary Fig S5).

Polymerization kinetics were then visualized using the mean intensities throughout all accepted condensates at any timepoint. Additionally, individual condensates were tracked throughout the timelapse using an existing tracking function [see https://site.physics.georgetown.edu/matlab/]. Tracking was terminated if coalescence events were detected based on the condensate volume. Further, averaged condensate images were extracted before and after the timelapse by averaging the images of condensates of identical radius.

### Actin kinetics theory

We employed a simplified model to recapitulate actin polymerization kinetics. The central assumptions are that the small aggregates, that are below the nucleus size, form and dissociate rapidly, but are more likely to dissociate. Once the nucleus size is exceeded, the rate for monomer addition is constant, i.e. independent of aggregate size. And finally, that the dissociation rate of a monomer from aggregates larger than and including the nucleus size is the same for all aggregate sizes (*17, 18*).

Taking a nucleus size of four actin monomers and utilizing previous notation (*18*) we constructed an iterative model for the molar concentration of filaments at time *t*, with *dt* = 1 s for simplicity,

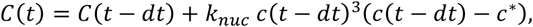

with molar concentration of filaments *C*, nucleation rate *k*_*nuc*_, molar concentration of G-actin *c* and critical concentration *c*^*^.

For the molar concentration of actin bound in filaments *F* we then found

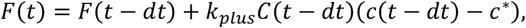

with association rate *k*_*plus*_. Note that the dissociation rate *k*_*minus*_ is captured in the critical concentration *c*^*^ = *k*_*minus*_/*k*_*plus*_.

We add an influx of G-actin to the system by adding a small fraction of G-actin in each time step. The data in Fig. 3G were calculated with *k*_*plus*_ = 10^6^ *M*^−1^*S*^−1^, *k*_*minus*_ = 1 *S*^−1^, i.e with *c*^*^ = 10^−6^*M, k*_*nuC*_ = 5 · 10^8^*M*^−3^*S*^−1^and *C*(*T* = 0) = 11 · 10^−6^*M*. The influx was set to 4.4 · 10^−10^*MS*^−1^.

The model is simplified and neglects the effects of phalloidin, which strongly affects the dissociation rate (*13*), as well as other effects, such as variation of ATP or ADP, and therefore is not expected to accurately reproduce the dynamics inside the condensate. The exact values of the nucleation kinetics are consequently likely not representative of the actual dynamics and have been chosen to roughly match experimental results while being comparable to literature values (*18, 37*). A closer look at the kinetics reveals that the linear regime occurs if the influx of G-actin is sufficient to sustain growth of existing filaments but not high enough to cause significant further nucleation of new filaments (which would induce non-linear growth). Moreover, changes in the kinetics affect the time scale of the process, but do not inhibit the emergence of the secondary regime, if an influx of G-actin is present, see Supplementary Fig. S6.

### Circular dichroism spectroscopy

For CD spectroscopy, PLPPR3 ICD was freshly purified in a reduced light absorbing composition of 20 mM phosphate buffer, 150 mM NaF and 1 mM DTT at pH 6.0. PLPPR3 ICD was diluted to 500 µL total volume with a final concentration of 5 µ M. CD spectroscopy was performed using a Jasco 5-720 spectroscope. First, a baseline scan against air was performed, second a scan with an empty cuvette with 0.1 cm light-path length and third with the buffer in the cuvette only. All measurements were performed at room temperature. Protein measurements were carried out with 200 µL PLPPR3 ICD in a cuvette previously washed with ddH_2_O, flushed with MeOH and dried with N_2_ gas. The following parameters were used for each run: sensitivity – standard 100 mdeg; start at 260 nm up to 195 nm (or until high-tension signal reached 800 V to prevent damage to photomultiplier). Data pitch at 0.1 nm; scanning mode in continuous; scanning speed 100 nm/min; response 1 s; band width 1 nm; accumulation 15 times to reduce measurement fluctuations. The accumulated curves were plotted against the wavelength.

To analyze the data and predict secondary structure elements, the baseline was subtracted from the measurements and the molar ellipcity calculated. The data were normalized to the number of aa residues of PLPPR3 ICD fusion construct (455) to yield the molar ellipcity/residue. To compare spectra more effectively, a data cut was introduced and the spectrum reduced to one data point per nanometer. The spectrum was compared to 35 standard spectra using the analysis software CDNN (Applied Photophysics Ltd.) to estimate secondary structure such as helices, anti and parallel β-strands, turns and coils. This gives an estimation of the total secondary structure or dynamic content.

### Mass spectrometry

#### Sample preparation

Protein samples dedicated for mass spectrometry analysis were diluted with RotiLoad (#K292.1, Roth) 1:4 (v/v), boiled 5 min at 95°C and frozen at -20°C. Samples were thawed, briefly boiled at 95°C and separated by 10% SDS-PAGE at 120 V for 2 hours (*38*). Proteins bands were visualized with Coomassie dye (#35081.01, Serva), gel regions containing visible bands cut out and stored at 4°C in 0.5 mL micro-centrifugal tubes containing 200 µL ddH_2_O. For identification and relative quantification of the proteins, gel pieces were subjected to tryptic digestion, as described in (*39*).

#### Liquid chromatography with tandem mass spectrometry analysis (LC−MS/MS)

Tryptic peptides were analyzed by LC-MS/MS using a Q Exactive Plus mass spectrometer (Thermo Fisher Scientific, Bremen, Germany). Peptide mixtures were fractionated by an Ultimate 3000 RSLCnano (Thermo Fisher Scientific) with a two-linear-column system. Digests were concentrated onto a 5 mm trapping guard column (PepMap C18, 300 μm x 5 μm, 100Å, Thermo Fisher Scientific). Then, samples were eluted from the analytical column a 75 µm i.d. × 250 mm nano LC column (Acclaim PepMap C18, 2 μm; 100 Å; Thermo Fisher Scientific). Separation was achieved by using a mobile phase from 0.1% formic acid (FA; Buffer A) to 80% acetonitrile (ACN) with 0.1% FA (Buffer B) and applying and 15 min active linear gradient from 3% to 53% of Buffer B at a flow rate of 300 nL/min. The Q Exactive instrument was operated in the data dependent mode to automatically switch between full scan MS and MS/MS acquisition. Survey full scan MS spectra (m/z 350–1650) were acquired in the Orbitrap with 70000 resolution (m/z 200) after 50 ms accumulation of ions to a 1 × 10^6^ target value. Dynamic exclusion was set to 10 s. The 10 most intense multiply charged ions (z ≥ 2) were sequentially isolated and fragmented by higher-energy collisional dissociation (HCD) with a maximal injection time of 120 ms, AGC 5 × 10^5^ and resolution 17500. Typical mass spectrometric conditions were as follows: spray voltage, 2.1 kV; no sheath and auxiliary gas flow; heated capillary temperature, 275°C; normalized HCD collision energy 27%. Additionally, the background ion m/z 445.1200 acted as lock mass.

#### Protein identification

Identification of the proteins was performed with Mascot software version 2.6.1. MS/MS spectra were searched with a precursor mass tolerance of 5 ppm, fragment tolerance of 0.02 Da, trypsin specificity with a maximum of one missed cleavage, cysteine carbamidomethylation set as fixed and methionine oxidation, protein N-acetylation as variable modification, against a self-created Fasta-database containing the PRG (PLPPR) modification.

### Crosslinking mass spectrometry

#### Sample preparation

The crosslinker DSS (#A39267, Thermo Fisher Scientific) was dissolved in DMSO to 50 mM (#A3672,0100, AppliChem) and diluted to 5 mM in a buffer composed of 20 mM HEPES, 150 mM NaCl at pH 6.0. Protein condensates were formed in 0.5 mL micro-centrifugal tubes (#0030108035, Eppendorf) with 20 µ M PLPPR3 ICD in the protein buffer (20 mM HEPES, 150 mM NaCl, 5 mM DTT at pH 6.0), 1:10 (v/v) of 10x F-actin buffer (1 M KCl, 0.1 M imidazole, 10 mM ATP, 20 mM MgCl_2_ at pH 7.4), 1.2 µM alpha-actin (#8101-01, Hypermol) and 5% (w/v) PEG8000 in a total volume of 20 µL. Where indicated, 1:10 (v/v) of 10x G-actin buffer (20 mM Tris-HCl, 4 mM ATP, 0.8 mM CaCl_2_, 0.1 mM DTT at pH 8.2) was substituted for F-actin buffer. Controls without condensates were obtained without adding PEG8000. In addition, actin controls were prepared with actin at 1.2 µM in F-actin buffer ±PEG, as well as actin at 10.2 µM actin in G-actin buffer and F-actin buffer-PEG. The mixture was incubated for 30 min at room temperature, before DSS was added 1:10 (v/v) to obtain a final concentration of 0.5 mM. The crosslinker was incubated for 30 min and quenched with Tris-HCl at pH 8.5 (final concentration: 50 mM) for 5 min.

#### Mass spectrometry preparation

The samples were reduced with DTT (final concentration 5 mM) for 30 min at 55°C and alkylated with chloroacetamide (#22788, Fluka, final concentration 40 mM) for 30 min in the dark. Roti-Load (#K292.1, Roth) was added 1:4 (v/v) and the samples boiled 5 min for 95°C. Samples were loaded on 10% SDS-gels (*38*) and the SDS-PAGE run at 120 V for 2 hours. Proteins were visualized with Coomassie dye (#35081.01, Serva), gel regions containing visible bands cut out and stored at 4°C in 0.5 mL micro-centrifugal tubes containing 200 µL ddH_2_O.

#### In-gel digestion

Gel bands were incubated with 50% acetonitrile (ACN) in 50 mM TEAB (pH 8.5) at 30°C for 10 min. After removing the washing buffer, 50 mM TEAB was added, incubated, and removed. Digestion was performed by incubating the gel pieces with 0.1 µg trypsin in 50 mM TEAB for 16 hours at 37 °C. The reaction was stopped by adding three volumes of 0.5% trifluoroacetic acid in ACN. The supernatant was dried along with peptides extracted from the gel pieces after an additional ACN wash.

#### Liquid chromatography and mass spectrometry

Peptides were resuspended in 1% ACN with 0.05% trifluoroacetic acid and injected into a Thermo Scientific™ Dionex™ UltiMate™ 3000 system. Separation was performed using a PepMap C-18 trap column (0.075 mm x 50 mm, 3 µm particle size, 100 Å pore size, Thermo Fisher Scientific), followed by an in-house packed C18 analytical column (Poroshell 120 EC-C18, 2.7 µm, Agilent Technologies). Peptides were separated at a flow rate of 250 nL/min using either a 117-min or 177-min gradient of increasing ACN concentration, and analyzed on an Orbitrap Fusion or Orbitrap Fusion Lumos mass spectrometer equipped with a FAIMS Pro™ device (Thermo Fisher Scientific), operated with Instrument Control Software versions 3.4 – 4.0. For MS1 scans, data were acquired in the Orbitrap with a resolution of 120,000, with advanced peak determination enabled. The MS1 acquisition settings were: m/z range of 375 – 1,600; standard AGC target; maximum injection time 50 ms. Precursors with charges of +4 to +8 were isolated within a 1.6 m/z window and dynamically excluded for 45 – 60 s. MS2 scans were performed in the Orbitrap at a resolution of 30,000 with the following parameters: automatic scan range; 200% AGC target; 100 ms maximum injection time; normalized collision energy (NCE) 30%. Data acquisition cycled between FAIMS compensation voltages of -40, -50, and -60 with a cycle time of 2 s per compensation voltage.

#### Database search

RAW data were analyzed using pLink2 (*40*) with the following parameters: DSS as the cross-linker, trypsin digestion allowing up to 3 missed cleavages, peptide mass range of 600 – 7,000 Da, and peptide length of 6 – 60 aa residues. Precursor mass tolerance was set to 10 ppm and fragment mass tolerance to 0.02 Da. Oxidation of methionine was included as variable, carbamidomethylation as a fixed modification. Inter- and intra-links were filtered separately to 1% false discovery rate (FDR) at the peptide-spectrum match (PSM) level. PLPPR-actin data were searched against the sequences of both proteins, common contaminants, and decoy sequences generated by reversing all entries. Crosslink spectrum match identifications were manually aggregated to peptide pairs.

#### Crosslinked peptide pair analysis and data visualization

All indentified crosslinked lysine/lysine peptide pairs were implemented into the evaluation. More specifically, each peptide pair combination (±PEG) of PLPPR3 ICD lysine-to-PLPPR3 ICD lysine, actin lysine-to-actin lysine, as well as PLPPR3 ICD lysine-to-actin lysine was counted, the sum calculated and compared to each other. Peptide pairs from 7 independent experiments were combined, which included PLPPR3 ICDs from different purification batches and actin. The differences in peptide pair identifications were visualized with Prism 9 and statistically evaluated using the Student’s t-test.

Total identified peptide pairs for PLPPR3 lysine/actin lysine combinations (+PEG) were substracted from identified peptide pairs (-PEG) to obtain the PEG-induced numeric difference. The data was visualized in a bubble chart with bubbles coloured in black indicative of peptide pairs ≥ 40, and bubbles in grey < 40 (numeric values are listed in Supplementary Fig. S10).

In 4 independent control experiments (without PLPPR3 ICD), peptide pair indentifications of actin lysine/actin lysine ± PEG were analyzed as mentioned earlier and specific peptide pairs i.e. K330/K328 were visualized (Supplementary Fig. S9 D). In addition, various peptide pairs were identified exclusively in control experiments with filamentous actin (10.2 µM actin; F-actin buffer; -PEG) compared to monomeric actin (10.2 µM actin; G-actin buffer; -PEG) i.e. K115/K193 (Supplementary Fig. S9 C).

### AlphaFold 3 models and PyMOL structures

AlphaFold 3 (*41*) was used to model interaction structures of PLPPR3 (uniport: Q7TPB0) with alpha-actin (uniport: P68032). Models were generated with the AlphaFold Server (https://golgi.sandbox.google.com/) with seed setting on auto. The models generated in JSON file format and protein structures available in the protein database (PDB) for F-actin (PDB-ID: 7Z7I) and WASP (PDB-ID: 2A3Z) were imported into the molecular graphics software PyMOL. For PLPPR3 ICD fragments modeled into actin (Fig. 4D), a broader amino acid sequencewas used for modeling. Whiledeletion variant Δ345 covers PLPPR3 aa 345-360, modeling was performed with aa 340-365. Likewise, for Δ399 covering PLPPR3 aa 399-417, models weregenerated with a broader sequencefrom aa 395-420. Following WH2-domain homology (Fig. 4C) and (*21*), only thecoremotif was deleted in Δ395 and Δ399. All molecular graphics representations were created with PyMOL 2.5.5 (Schrödinger, LLC.) utilizing the ray command at high resolution (ray 3000, 3000).

## Supplementary Figures

**Fig. S1.**
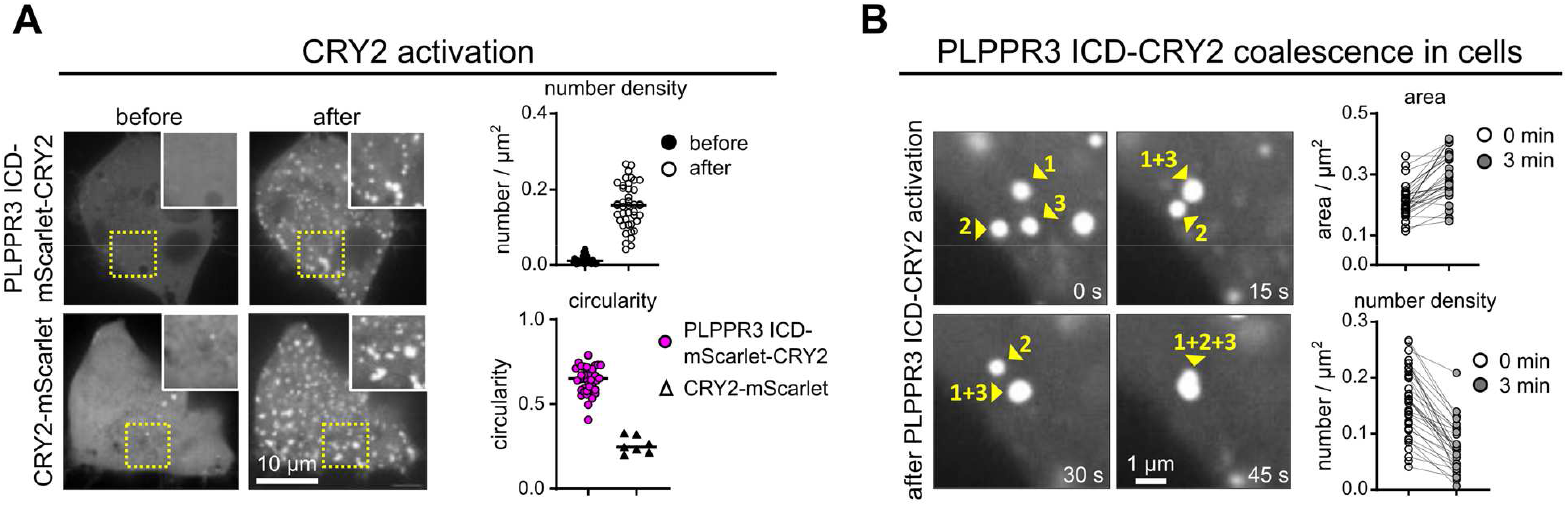
Additional data for PLPPR3 ICD-CRY2 activation in cells. **(A)** Number density of PLPPR3 ICD-mScarlet-CRY2 puncta before and after activation. Circularity of PLPPR3 ICD-mScarlet-CRY2 puncta compared to CRY2-mScarlet control after activation. **(B)** Representative images of coalescence events. The area of condensates increases over time, whereas the number density decreases, indicating coalescence. All data were analyzed in ImageJ.

**Fig. S2.**
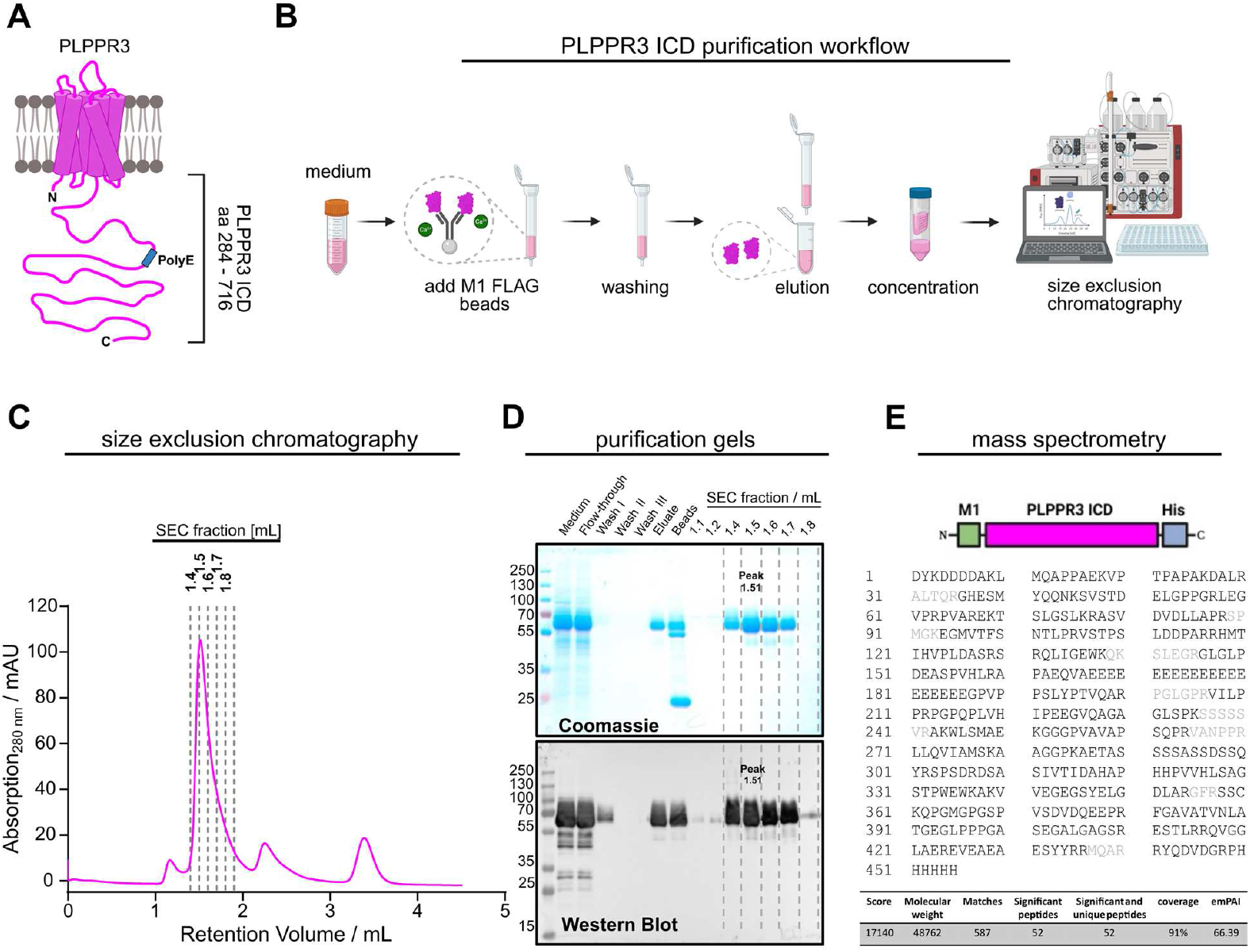
Purification and analysis of PLPPR3 intracellular domain (ICD) from Expi293F cells. **(A)** PLPPR3 consists of transmembrane regions (amino acid residues 1-283) and a long intracellular domain (ICD) corresponding to amino acids residues 284 to 716. **(B)** The construct HA-M1-PLPPR3 ICD-His was overexpressed for 96 hours in Expi293F cells. PLPPR3 ICD was captured from the cellular supernatant via Ca^2+^ dependent M1 Flag beads, washed and eluted via competitive M1 Flag peptide and EDTA. Protein was purified via size exclusion chromatography, concentrated and stored at -80°C. **(C)** Size exclusion chromatography with homogeneous protein peak at 1.51 mL. The fractions at 1.4 mL – 1.8 mL were analyzed by SDS-PAGE stained by Coomassie blue (**D, top**) and by western blot (**D, bottom**) with a specific α-PLPPR3 antibody. **(E)** PLPPR3 ICD was analyzed by mass spectrometry. Peptides marked in black were detected, while peptides in grey were not. The overall coverage rate was > 90%.

**Fig. S3.**
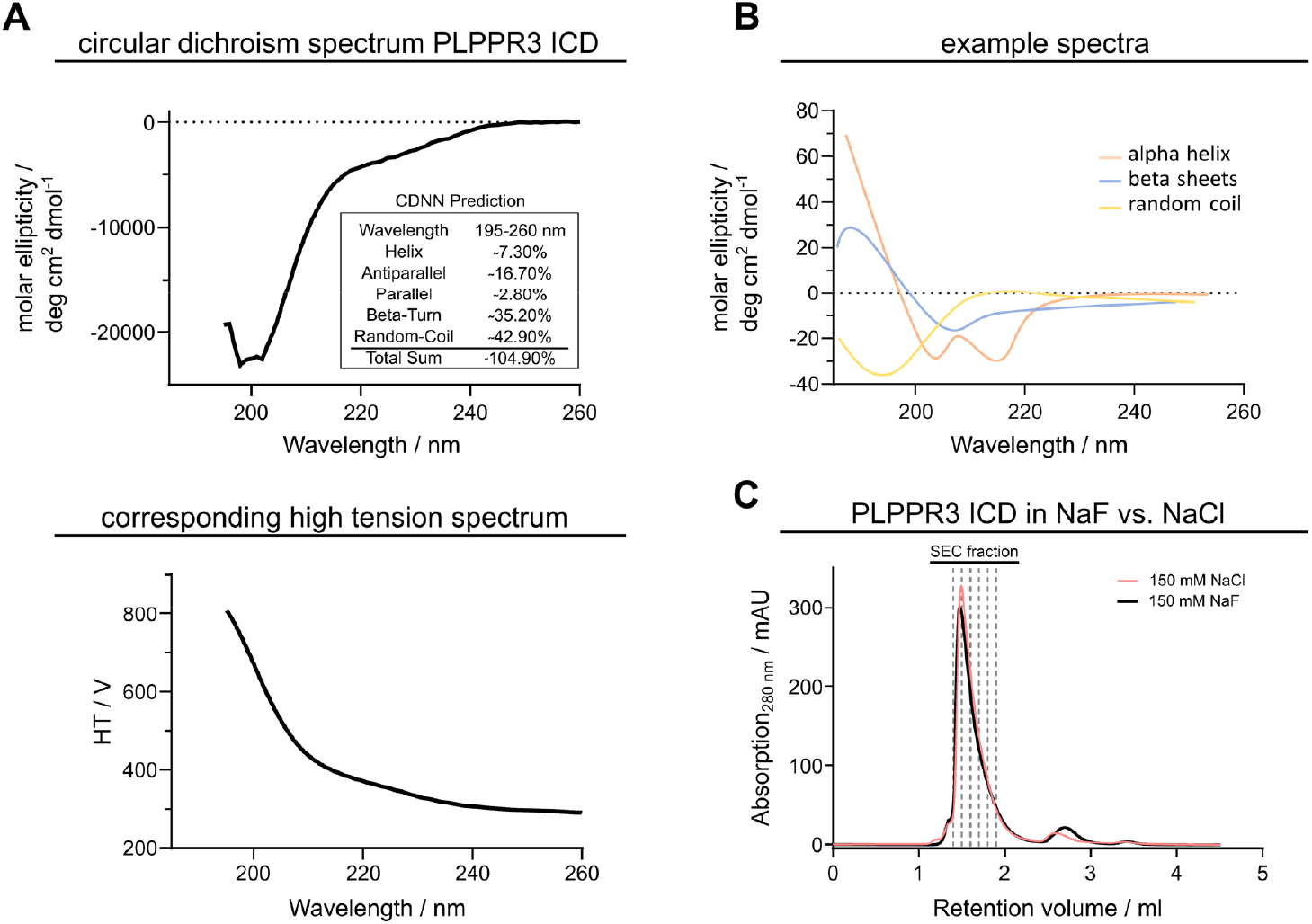
PLPPR3 intracellular domain (ICD) shows characteristics of an unstructured protein in circular dichroism spectroscopy. **(A, top)** Circular dichroism (CD) spectrum of purified PLPPR3 ICD in 20 mM phosphate buffer, 150 mM NaF, 1 mM DTT at pH 6.0 indicates that large parts of the proteins are unstructured. CDNN analysis software predicts roughly 78% in a conformationally dynamic state. **(A, bottom)** corresponding high tension spectrum as a measurement of CD spectrum quality. Especially in far-UV CD the light is absorbed more strongly at lower wavelengths, which requires the detector to increase voltage to maintain a stable signal (*42*). **(B)** Representative example spectra for different secondary structure elements; adapted from (*42*). **(C)** To reduce light absorption of the sample during CD measurements, PLPPR3 ICD was purified in an alternative buffer system. PLPPR3 ICD was purified in 20 mM HEPES, 150 mM NaCl, 5 mM DTT at pH 6.0 (red line) and compared to purified ICD in 20 mM phosphate buffer, 150 mM NaF, 1 mM DTT at pH 6.0 (black line). The protein was stable in the alternative buffer system.

**Fig. S4.**
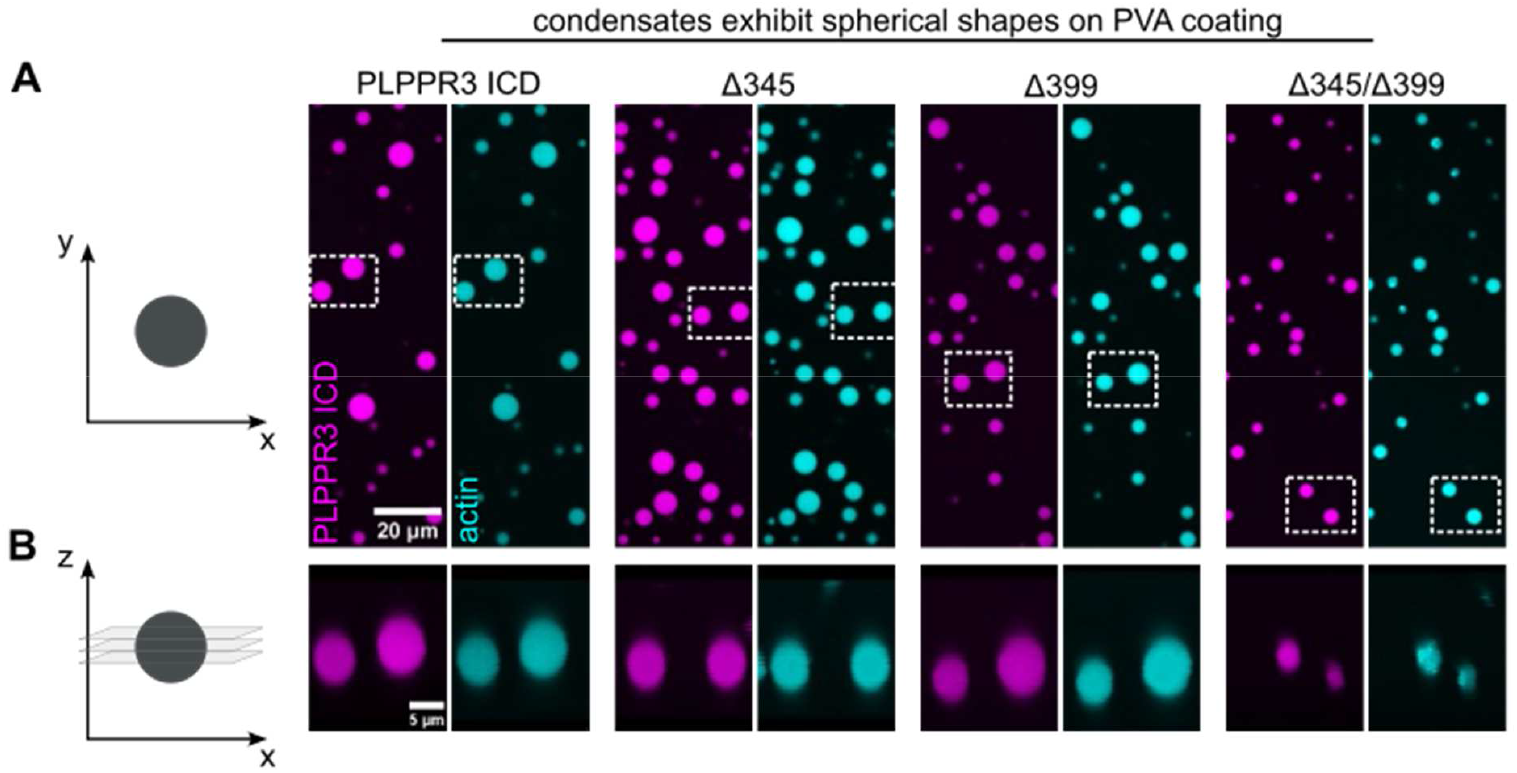
Condensates exhibit spherical shapes on polyvinyl alcohol (PVA)-coated coverslips. PLPPR3 intracellular domain (ICD) and deletion variants form condensates with 40 µM ICD concentration and 5% (w/v). Actin partitions into all condensates, see Fig. 5A for quantification. Imaging was performed 1 hour after addition of PEG. **(A)** Top view on PLPPR3 ICD and actin condensates. **(B)** Side view displaying spherical shapes on PVA-coated coverslips. Stacks were generated with ImageJ. All condensates exhibit non-wetting conditions.

**Fig. S5.**
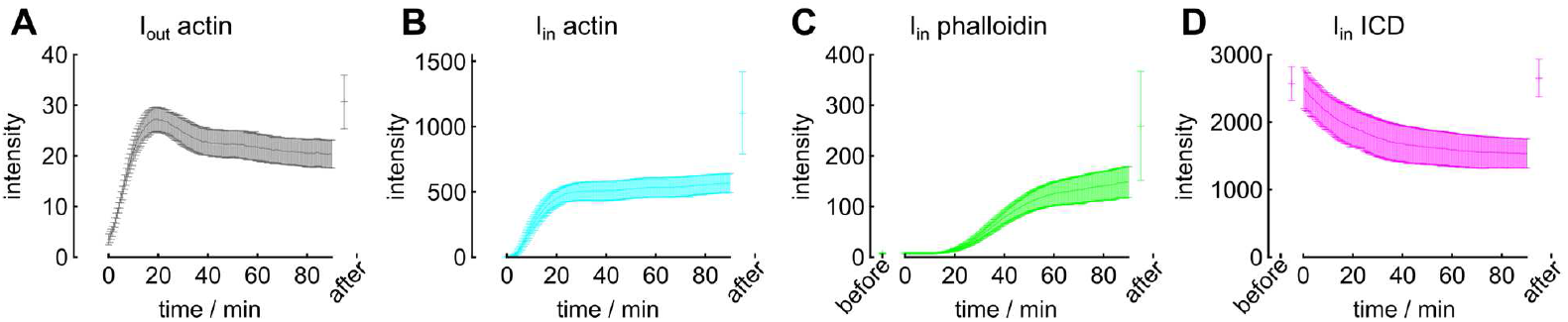
Non-normalized intensity profiles of data from Fig. 3F reveal the extent of photobleaching. Data show the mean with standard deviation for samples before, during and after timelapse imaging for A) *I*_*out,actin*_, B) *I*_*in,actin*_, C) *I*_*in,phalloidin*_ and D) *I*_*in,ICD*_. Note, that data before and after timelapse imaging are collected in different fields of view than those collected during the timelapse.

**Fig. S6.**
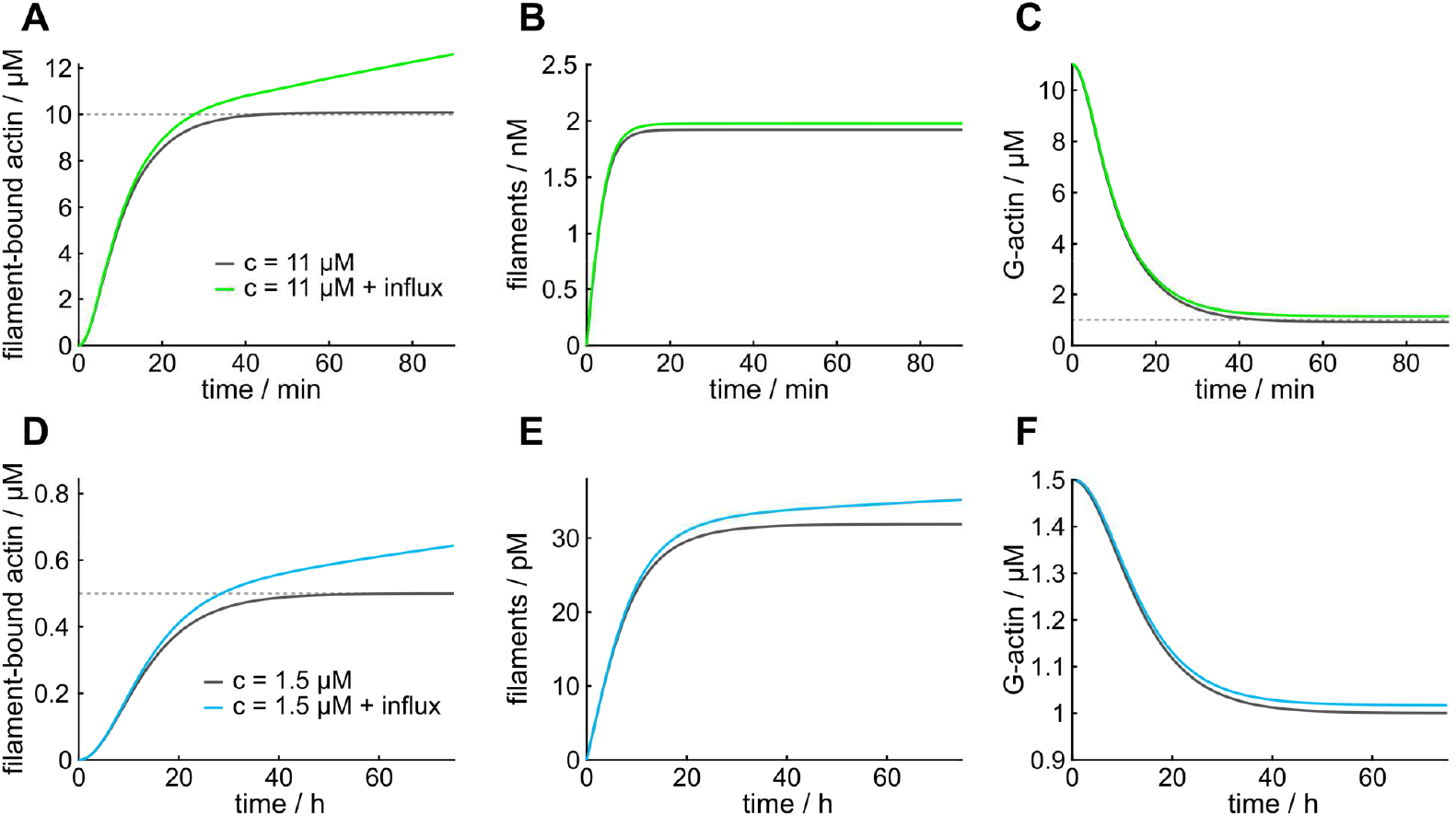
Theoretical actin polymerization kinetics. **(A)** Molar concentration of filament-bound actin over time. The dashed line shows the total concentration minus *c*^*^. **(B)** Molar concentration of filaments over time. **(C)** Molar concentration of G-actin over time. The dashed line shows *C*^*^ = 1 *μM*. Actin polymerization model in panels (A-C) calculated with *k*_*plus*_ = 10^6^ *M*^−1^*S*^−1^, *k*_*minus*_ = 1 *S*^−1^, *k*_*nuc*_= 5 · 10^8^*M*^−3^*S*^−1^, and *c*(*T* = 0) = 11 · 10^−6^*M*. The influx was set to 4.4 · 10^−10^*MS*^−1^. **(D)** to **(F)** The same quantities calculated with minimal oversaturation in the system results in the same functional form but with significantly slower dynamics. Calculated with *k*_*plus*_ = 10^6^ *M*^−1^*S*^−1^, *k*_*minus*_ = 1 *S*^−1^, *k*_*nuc*_ = 5 · 10^8^*M*^−3^*S*^−1^, and *c*(*T* = 0) = 1.5 · 10^−6^*M*. The influx was set to 6 · 10^−13^*MS*^−1^.

**Fig. S7.**
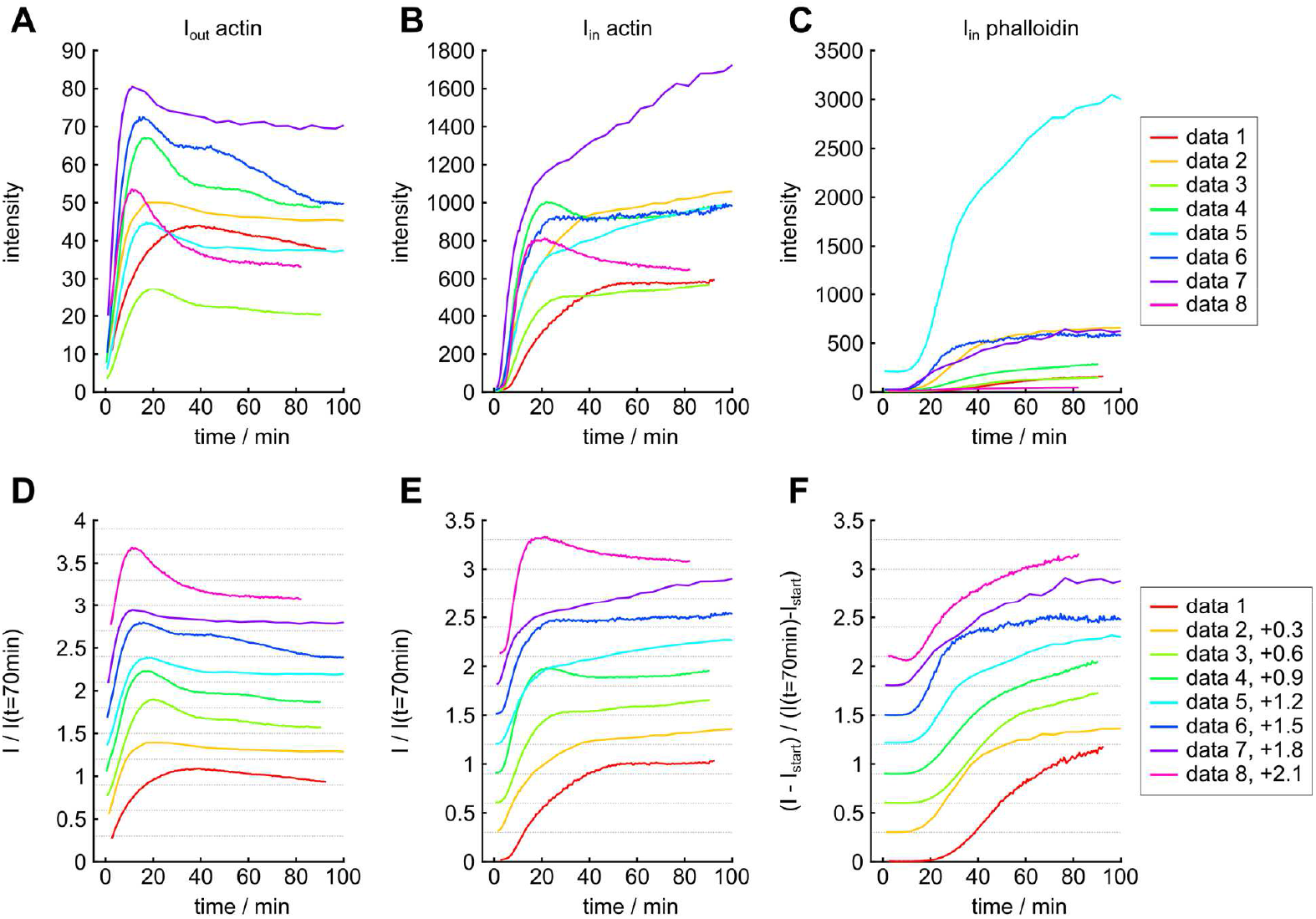
Intensity profiles of 8 independent actin polymerization assays. Unnormalized intensity profiles for I_out_ actin **(A)**, I_in_ actin **(B)** and I_in_ phalloidin **(C)**. All data aligned in time by their respective time of actin arriving outside the condensate t_act,out_, extracted from a linear fit to I_out_ actin. Panels **(D-F)** show the same data rescaled by their respective mean value around t=70 min. An offset of 0.3 (dashed lines) is added iteratively to the data for better comparability.

**Fig. S8.**
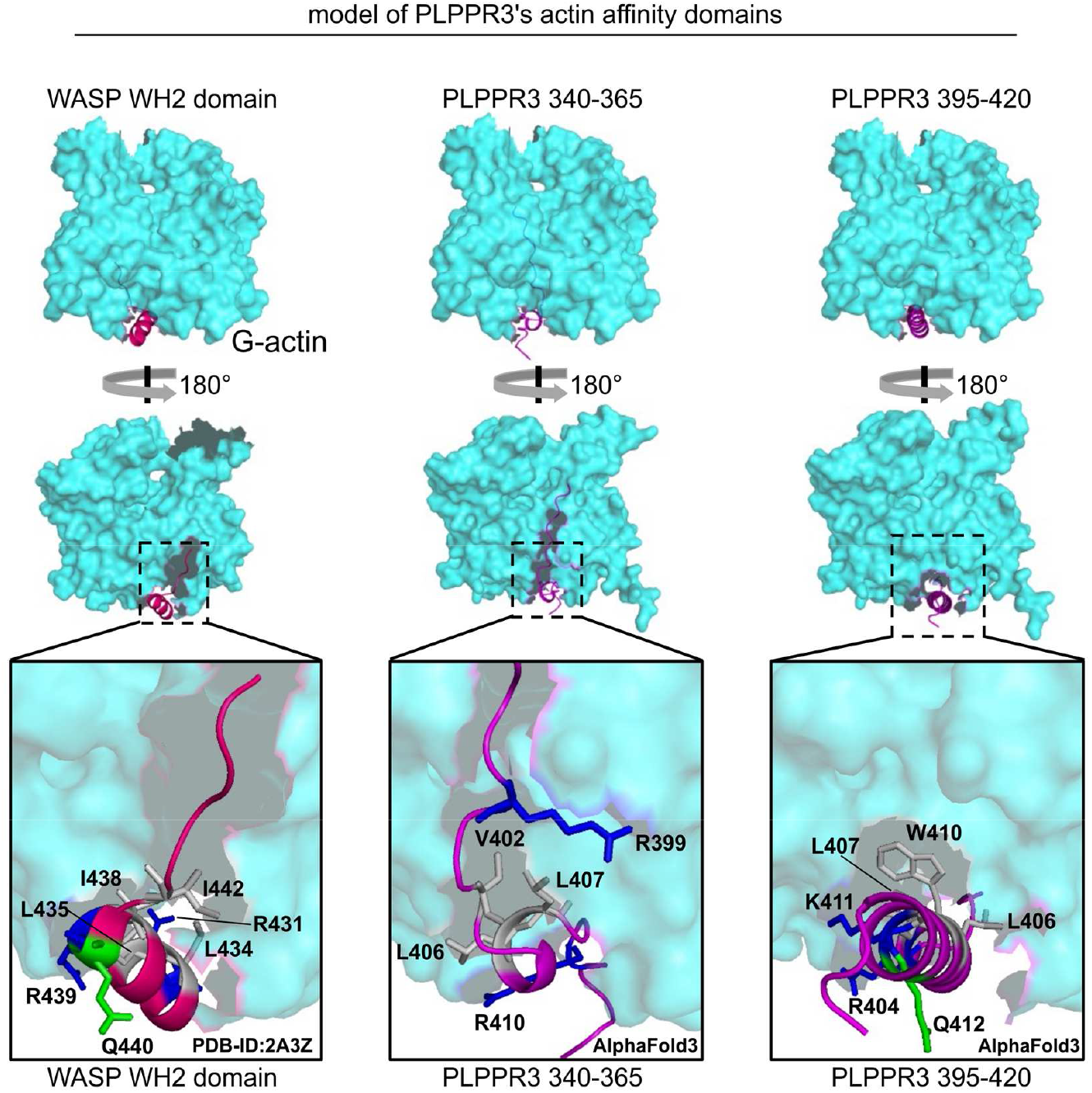
WH2 actin binding domains of WASP and WH2-like actin affinity domains of PLPPR3. **(Left)** G-actin model (PDB-ID: 2A3Z) with the WASP WH2 domain in the ABC. Core WH2 amino acid residues for the actin interaction are marked using the same color code shown in Fig. 4D (adapted from (*21*)). **(Middle)** AlphaFold3 model of G-actin with PLPPR3 aa 340-365 in the ABC forming a helix-like structure. **(Right)** AlphaFold3 model of G-actin with PLPPR3 aa 395-420 creating a helix-forming structure. Note that both PLPPR3 340-365 and PLPPR3 395-420 contain WH2-like homology (Fig. 4C).

**Fig. S9.**
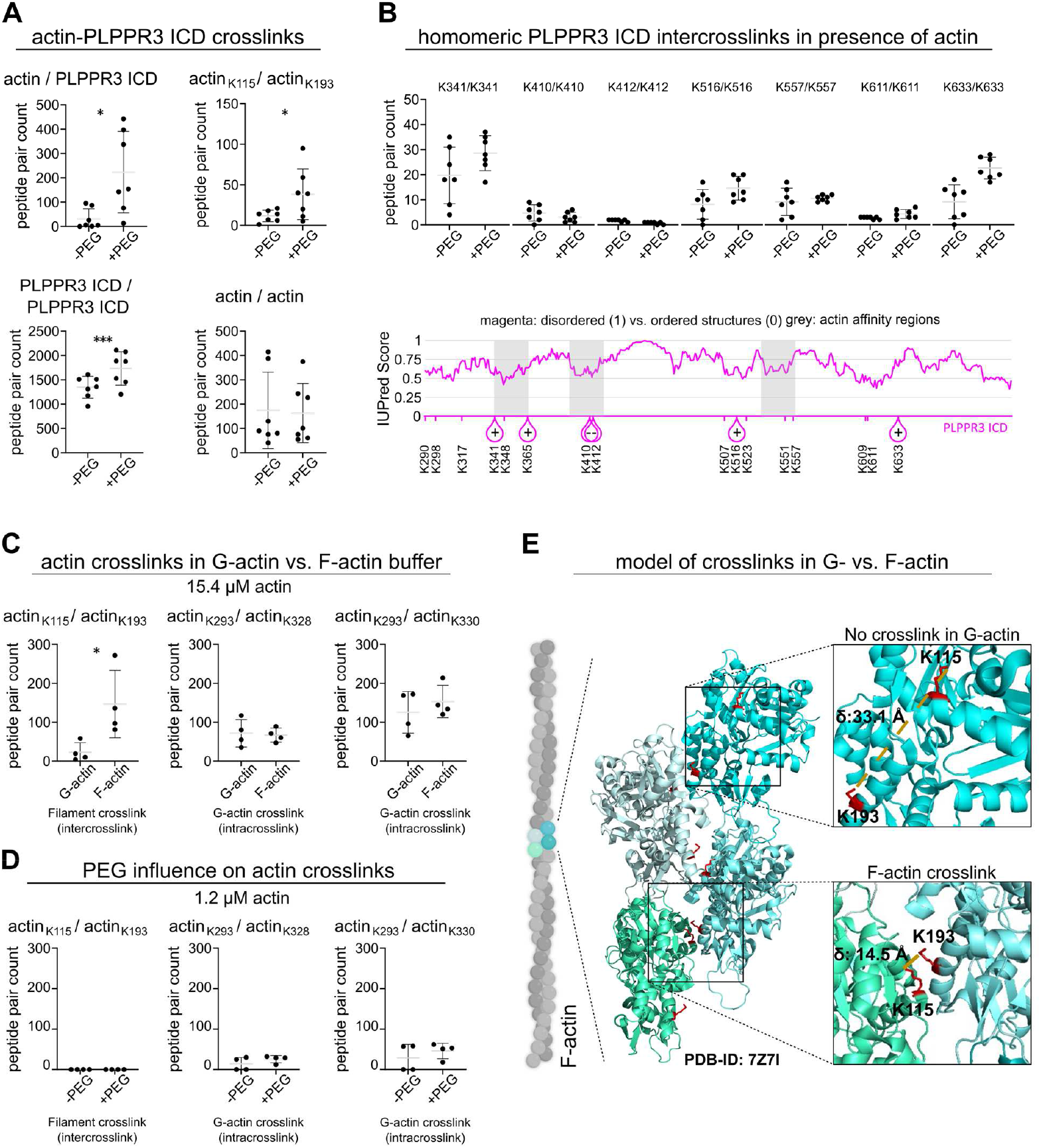
Controls for crosslinking experiments. **(A)** Crosslinks in samples containing 1.2 µM actin and 20 µM ICD under the same conditions as the polymerization assay, but without phalloidin, with and without 5% (w/v) PEG to induce phase separation. Panels show crosslinks between actin and ICD, the F-actin specific crosslink K115 to K193, crosslinks within ICD proteins and within actin proteins. **(B)** Homomeric inter-crosslinks of PLPPR3 ICD in the presence of actin. Lower panel: IUPred structure prediction (https://iupred3.elte.hu/) of PLPPR3 ICD overlain with locations of homomeric crosslinks. **(C)** Actin crosslinks in G- and F-actin buffer without ICD at a G-actin concentration of 15.4 µM. The F-actin specific crosslink K115 to K193 increases in F-actin buffer. **(D)** Impact of 5% PEG on actin under polymerization assay conditions without PLPPR3 ICD. **(E)** Visual representation of the F-actin specific crosslink K115 to K193.

**Fig. S10.**
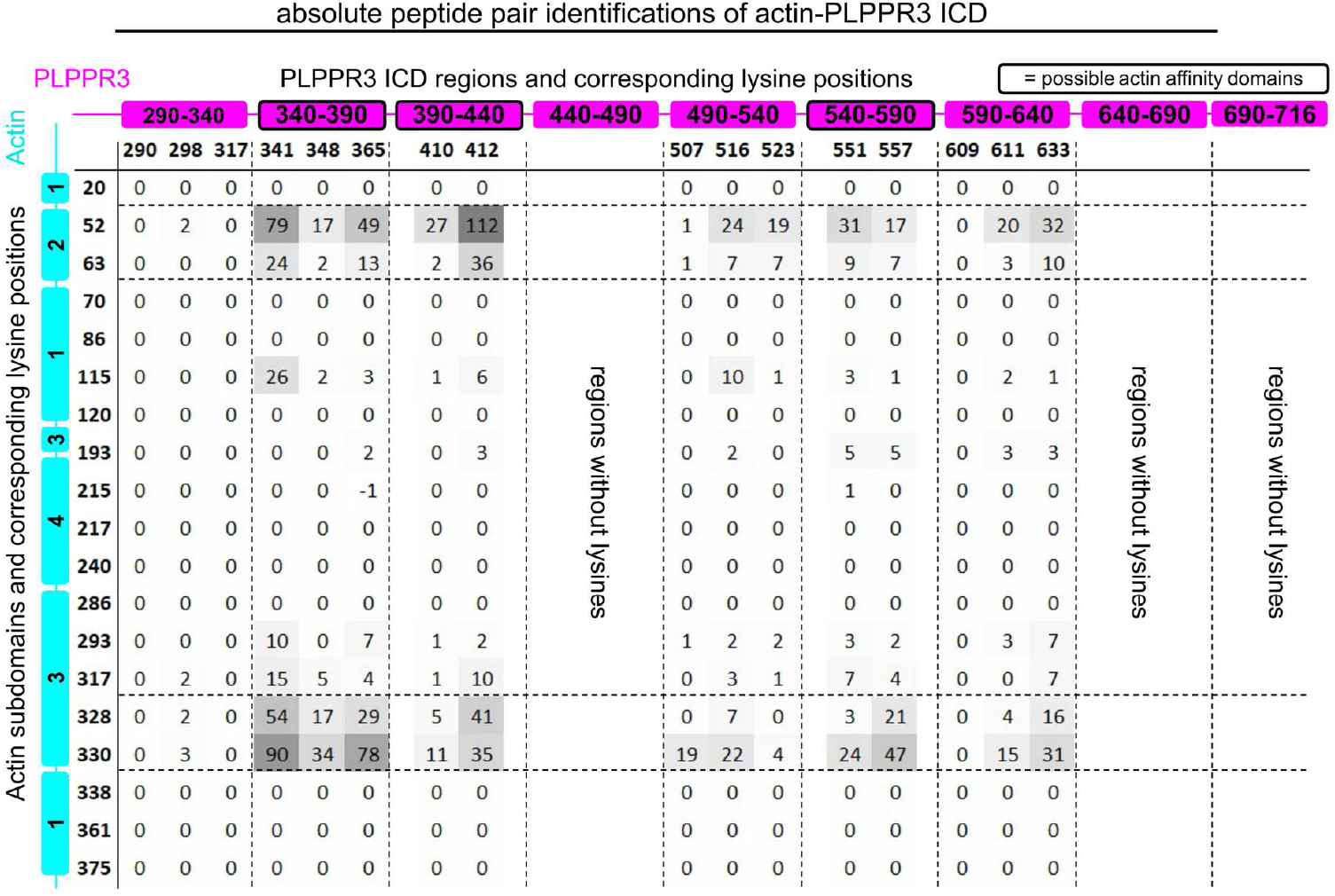
Crosslinking data from mass spectrometry (XL-MS) (tabular format). Identified crosslinked peptide pairs between PLPPR3 ICD and actin under PEG-induced condensate conditions minus crosslinked peptide pairs from conditions without PEG. The table shows the relative abundance of PLPPR3 ICD-actin crosslinked peptide pairs (N=7). Gray shades highlight high occurrence.

**Fig. S11.**
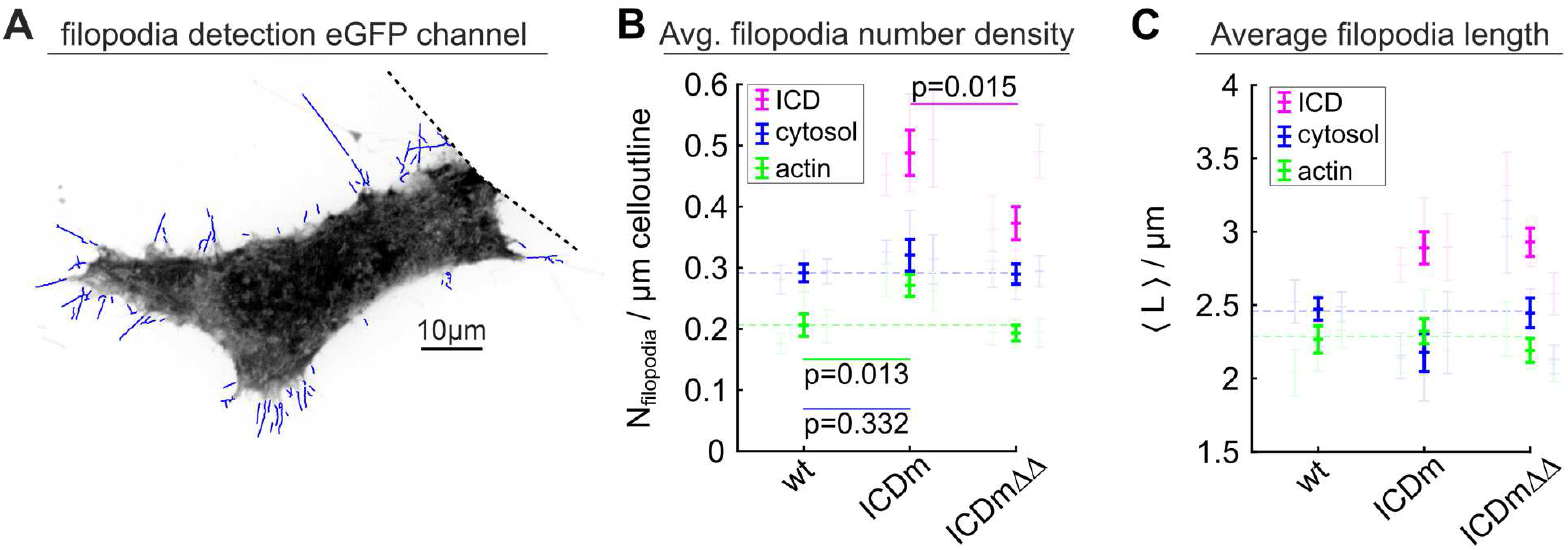
Filopodia analysis including cytosol stain via eGFP transfection. **(A)** Representative cytosolic eGFP staining overlayed with detected filopodia (blue). Same cell as in Fig. 5C. **(B)** Average filopodia number density including the cytosol stain. Highlighted data show the overall mean and standard error, data in the background show independent experiments. **(C)** Average filopodia length.

